# *Drosophila* embryo cellularization is modulated by the viscoelastic dynamics of cortical-membrane interactions

**DOI:** 10.1101/2025.02.07.637185

**Authors:** Kyle Stark, Mayte Bonilla Quintana, Anna Marie Sokac, Padmini Rangamani

## Abstract

The generation of an epithelial sheet transforms fruit fly embryos from a single syncytial cell directly into a tissue. For this to happen, the apical microvillus membrane is pulled between peripherally anchored nuclei in a process known as furrow invagination. Experimental measurements of furrow invagination velocities have shown that the rate of invagination undergoes slow-to-fast and fast-to-stalled velocity transitions during the formation of individual cells. The causes of such changes are due to multiple intersecting mechanisms and molecular components, including motor proteins, microtubules, and F-actin. In this work, we develop a continuum model to describe the dynamics of furrow invagination. Our model is constrained by previously published experimental data and considers the roles of cytoskeletal forces, cytoplasmic drag, motor protein forces, and membrane tension. We find that the viscous forces produced by the cytoskeleton sliding beneath the plasma membrane dictates furrow velocity. We propose that the slow phase is slow because there is a high density of microvilli, which increases the number of viscous contact points between the plasma membrane and the underlying cytoskeleton. This in turn, results in a higher resistance to furrow invagination. We predict that the fast phase may benefit from fewer cytoskeleton-to-plasma membrane contact points, thus reducing viscous forces and promoting the slow-to-fast switch. Then, we use perturbation and loss-of-function simulations to show that microvillus and sub-apical membrane reservoirs are vital to setting furrow invagination dynamics. This work demonstrates how coupling between the cytoskeleton, the plasma membrane, and distinct membrane reservoirs affects the plasticity and dynamics of cellularization.

## 3 Introduction

*Drosophila* embryos are directly transformed from a single syncytial cell into a tissue by deploying a novel form of cellular division, called cellularization [1, 2]. To cellularize, the embryo simultaneously divides its apical surface into individual mononucleate cells by pulling invaginating furrows between each of the ∼ 6000 peripherally-located nuclei [2] (Fig. 1A, B). The apparent surface area of the membrane is increased by unwrinkling the microvillus membrane located on the apical surface of the embryo [3]. This strategy is a critical event in the fly, as disruption leads to developmental failures [4–6]. Similar furrow invaginations are necessary for the proper function of striated muscle and mitochondria, by forming transverse-tubules (t-tubules) and cristae, respectively. Thus studying these structures in the fly provides an opportunity for understanding the mammalian counterparts [7–11].

**Figure 1:**
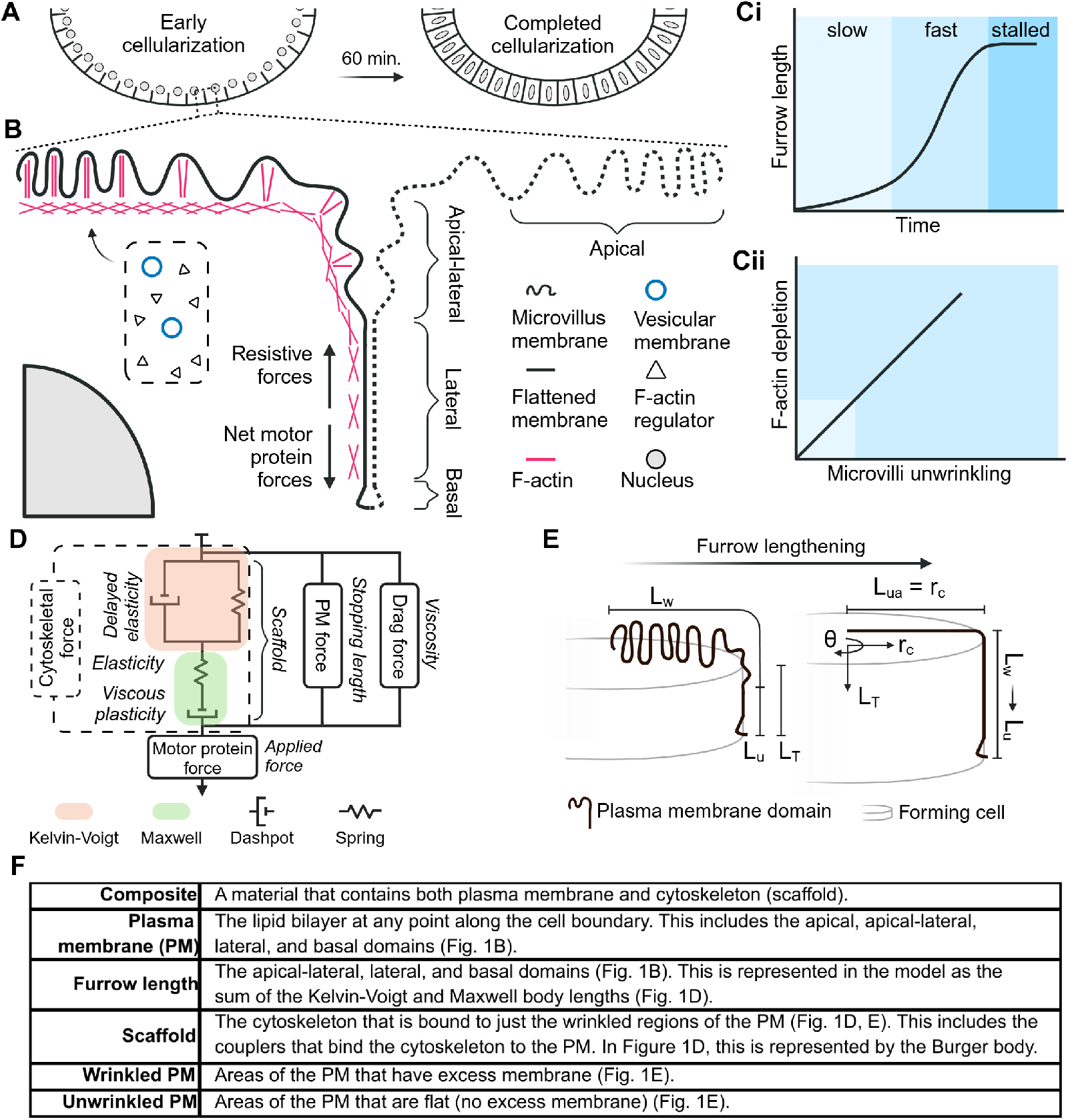
A computational model of furrow invagination. Figure 1. : (A) Cellularization generates the primary epithelium of the *Drosophila* embryo. (B) The force balance is evaluated at the tip of the plasma membrane-cytoskeleton composite (solid plasma membrane (PM), basal region); the dotted PM of the neighbor cell shows the microscopic scale of cell periodicity and is not a part of the stress balance in the model. The microvilli on the surface of the embryo unwrinkle to provide membrane surface area for furrow invagination over ∼ 60 min. Cytoskeleton regulators and vesicle exocytosis promote the assembly of microvilli. Vesicles exocytose apically in the slow phase (first 30 - 40 min). (C) A schematic of experimental observations from [3, 21]. Furrow invagination velocity switches from slow to fast and fast to stalled (Ci). Experiments show that microvilli unwrinkling and F-actin depletion from microvilli are positively correlated (Cii). (D) A schematic representation of the stress balance that leads to the governing equations in the model (Eq. 8). (E) Conservation of membrane area; here *L*_*ua*_, *L*_*u*_, *L*_*w*_, *L*_*T*_, *r*_*c*_ are unwrinkled apical length, unwrinkled lateral length, wrinkled length, total length, and cell radius, respectively. Wrinkled area in the cell membrane is only present on the apical surface and is converted to unwrinkled length as the furrow length increases. Unwrinkled apical length is a constant value and accounts for the end-state when the apical surface becomes unwrinkled. (F) A table of key terms used throughout the paper.

The invagination of the *Drosophila* furrow is fundamentally a cell mechanics problem because it is a time-dependent change in cellular shape that is driven by mesoscale forces. We hypothesize that furrow formation is governed by the plasma membrane (PM), cytoskeleton, cytoplasm, and a motor protein driving force [12–15] (Fig. 1B). Here, the cytoskeleton and PM form a composite that defines the material properties and shape of the furrow. The furrow is pulled intracellularly by energy-consuming motor proteins [16]. As this happens, the cytoplasm and PM-bound cytoskeleton offer a viscoelastic resistance on the furrow [12–15].

Besides forming a composite with the cytoskeleton, the plasma membrane is crucial for furrow invagination. Specifically, the microvillus membrane on the apical surface of the cell stores excess membrane necessary for the furrow to reach a final length of 35 µm [3, 6, 17]. Upon reaching the final length, the irregular microvillus surface becomes smooth, indicating that this external membrane reservoir may limit furrow length [3, 6]. However, only about 50% of the necessary membrane is stored in the microvilli at the start of cellularization, which is sufficient to support 10-15 µm of furrow invagination [3, 17]. The additional membrane required for successful furrow invagination is exocytosed from the Golgi to the microvilli during the first half of cellurization (30-40 minutes), which allows for continued furrow invagination in the latter half of cellularization [3, 17]. Curiously, when the delivery of vesicles to the microvilli stops, the furrow velocity jumps from ∼ 3.5 nm/sec to ∼ 24 nm/sec [3]. Additionally, the elastic and viscous coefficients of the apical surface drop by ∼ 5x and ∼ 4x, respectively [13]. These observations allow us to divide furrow invagination into distinct slow, fast, and stalled phases (Fig. 1C). Finally, mechanical measurements of the apical PM-bound cytoskeleton reveal that it behaves as a fluid-type viscoelastic body and that F-actin correlates positively with microvilli density (Fig. 1C) [6, 13, 14].

The above experimental observations indicate that the dynamics of furrow invagination are affected by a coupling between the viscoelastic properties of the PM-bound cytoskeleton, and the kinetics of vesicle transport to the PM. We seek to understand possible mechanisms that govern (1) the slow-to-fast velocity shift, (2) the fast-to-stalled velocity shift, and (3) how dynamic regulation of membrane area influences furrow behavior. Our model leverages previous modeling efforts by D’Angelo *et al*. [13] and Doubrovinski *et al*. [14], which demonstrated that the apical surface has a fluid-like behavior when measured at discrete instances in time and that a shift in material properties occurs at the slow-to-fast transition. Furthermore, we incorporated the idea of a limitation in membrane area as originally proposed by Sommi *et al*. [16]. However, the model proposed by Sommi *et al*. [16] used separate sets of equations to govern the slow and fast phases. While each of these models interrogated specific, discontinuous aspects of furrow invagination, we combined membrane area, membrane reservoirs, and the PM-cytoskeleton composite to replicate invagination dynamics continuously in time and with a single set of equations.

Our findings propose that softening of the PM-cytoskeleton composite produces the slow-to-fast shift. We predict that membrane area restriction is a possible mechanism of stopping furrow invagination and hypothesize that membrane is added through exocytosis of vesicles from the Golgi and a secondary, internal sub-apical compartment. Moreover, we investigate how the transport of membrane and cytoskeletal elements [16, 18–20] can be coupled to change the material properties of the furrow. By mimicking *in vivo* perturbation and loss-of-function experiments, we constrained our model to experimental measurements from the literature and predicted that insertion of cytoskeletal elements at the apical-lateral region of the furrow softens the cortex to produce the slow-to-fast shift in the rate of furrow invagination. In summary, with a single set of equations, our model was able to replicate the key experimentally observed features of furrow invagination including the membrane area dynamics, the slow-to-fast and fast-to-stalled velocity shifts, and isolated a plausible mechanism underpinning the sudden softening of the material properties.

## 4 Results

The length of the furrow is governed by a force balance at the basal tip of the furrow as shown in Figures 1B and D. The force balance at the tip is given by

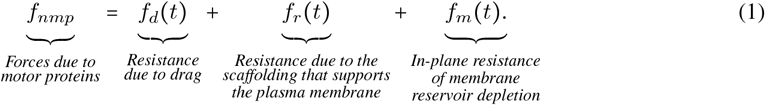

Here, *f* represents the force per circumferential length of the cell being formed. The net motor protein force, *f*_*nmp*_, drives invagination by pulling the furrow inward [2, 16] and all other forces resist invagination. The drag force, *f*_*d*_ (Eq. 18), accounts for the resistance imposed by a viscous medium on the lengthening furrow. Resistance due to the PM-bound cytoskeleton (referred to as “scaffolding” for convenience) that supports the PM, *f*_*r*_ (Eq. 12), results from cytoskeletal network deformation. The portion of the scaffold that stabilizes the wrinkled membrane reservoir deforms as the furrow lengthens [3, 4, 6]. In-plane membrane resistance, *f*_*m*_ (Eq. 19), is expected to rise rapidly when the wrinkled membrane reservoir is depleted.

In sections 4.1, 4.2, and 4.3 we develop and simulate a system of ODEs in time, to discover the minimal mechanics governing furrow invagination dynamics. Then, in sections 4.4 and 4.5, we test possible causes of the material softening that were identified in sections 4.2 and 4.3. Finally, in section 4.6, we explore the applicability of our model by recapitulating perturbation and loss-of-function experiments. Detailed model equations are given in the Methods section and parameters in Tables S1 and S2. In the Results section, we have reproduced some of these equations for readability.

### 4.1 Resistive forces of furrow invagination are captured by a submerged viscoelastic body

We first propose that one of the resistive force felt by the furrow is from the scaffold, *f*_*r*_, and that it behaves in a viscoelastic manner (Fig. 1D). Predictive models of PM-bound cytoskeleton support this modeling decision [12–14, 22, 23]. Physically, our viscoelastic model represents the entire scaffold along the PM (Fig. 1B) – not individual microvilli or molecular details – because that is the spatial resolution of our experimental data (Eqs. 13-15) [2, 3, 13, 14]. Our specific model uses a Kelvin-Voigt (KV) body in series with a Maxwell (MW) dashpot because they collectively capture the delayed elasticity and viscous plasticity identified by D’Angelo *et al*. [13] and Doubrovinski *et al*. [14] (see Fig. 1D for a schematic representation and Findley *et al*. [24] as an excellent source on viscoelasticity). Briefly, a Kelvin-Voigt body is elastic (solid-like) because it returns to its initial length after any external loading that caused it to stretch is removed. However, the elasticity is “delayed” because of energy loss through the parallel dashpot. Biologically, this delay can be attributed to friction between molecular constituents. While the Maxwell dashpot also experiences molecular friction, it does not recover its initial length (also referred to as “permanent” or “plastic” deformation) because it lacks the restoring force of a parallel spring, as is found in the KV body. The final element in our viscoelastic body is a series spring, which captures the finding from Herant *et al*. [23] that microvilli can undergo an initial, frictionless “slack” when stretched (Fig. 1D). Collectively, this configuration of elements is referred to as a Burger body (Section 6.2.2) and contains elasticity, viscosity, delayed elasticity, plasticity, instantaneous recovery, and delayed recovery [24]. From this starting point, we then identify which features of the Burger body best describe furrow invagination. Each feature is labeled in Figure 1D, except for recovery features since the furrow only lengthens during cellularization [3], i.e., there is always an applied load by the motor proteins. Finally, for a more realistic viscoelastic representation we add viscous drag, *f*_*d*_, to simulate submerging the *in silico* furrow in the cytoplasm (Eq. 18, Fig. 1D) [25].

### 4.2. The slow-to-fast transition requires time-dependent changes to the material properties that resist invagination

We test if the simplest case – a system with parameters that are constant in time – can produce the slow-to-fast velocity shift of the invaginating furrow. We analyzed the dynamics of the Burger body under a constant applied force (i.e., *f*_*r*_ (*t*) in Eq. 12 is *f*_0_). The body is composed of three series components: the Kelvin-Voigt body, the Maxwell spring, and the Maxwell dashpot (Fig. 1D). To find the length dynamics of the system, we simply add the length dynamics of each of its uncoupled components (Fig. 2A, B). Therefore, the total length is *L*_*T*_ = *L*_*rk*_ + *L*_*rms*_ + *L*_*rmd*_, where *L*_*rk*_, *L*_*rms*_, and *L*_*rmd*_ are the lengths of the Kelvin-Voigt body, the Maxwell spring, and the Maxwell dashpot, respectively. Rearranging the equation for the force felt by the KV body (Eq. 13), we observed that the Kelvin-Voigt velocity decays as a function of its length, 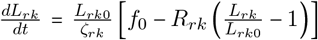. Here, the first term, 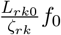, is the maximum rate of extension of the KV body, while the second term, 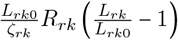, autoinhibits the rate of extension due to elastic strain. The Maxwell spring length (Eq. 14) has no time-dependence as, 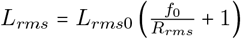, and therefore, *L*_*rms*_ changes instantaneously 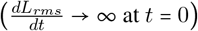 to a new steady-state length 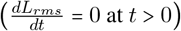. The Maxwell dashpot velocity (Eq. 15) increases at a constant rate, 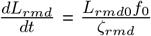. Therefore, the velocity of the total Burger body slows monotonically as 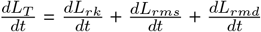

**Figure 2:**
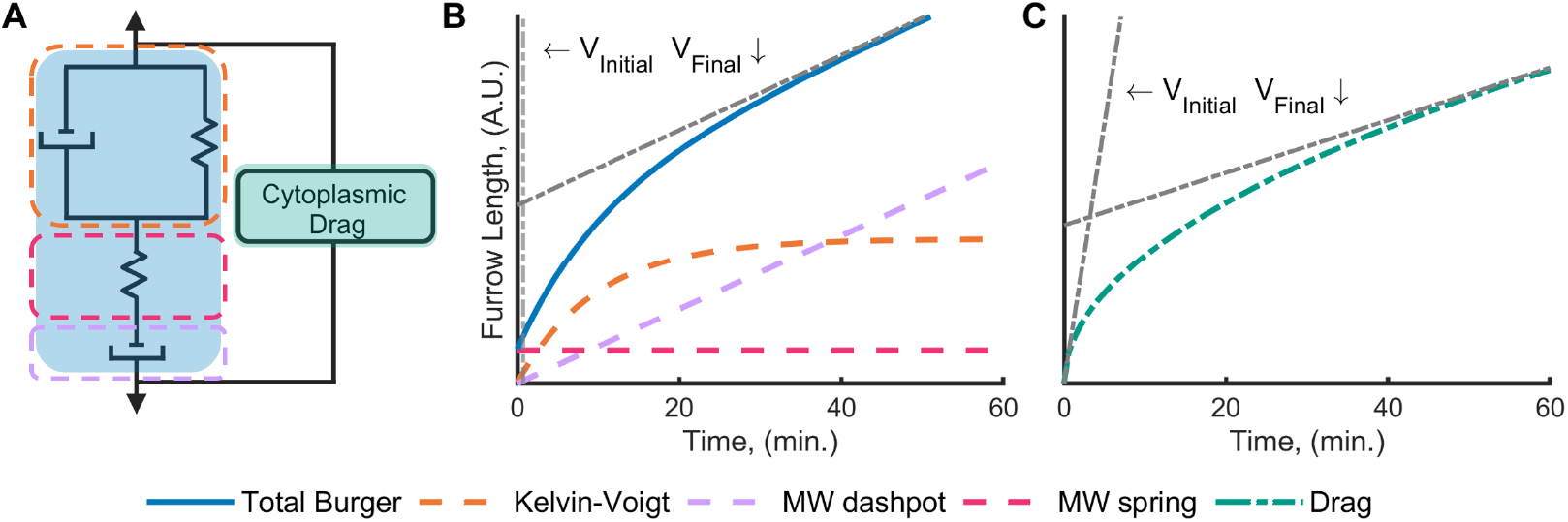
Viscoelastic coefficient softening is necessary to achieve a slow-to-fast velocity shift. (A) A cartoon representation of the cytoskeletal (blue shading) and cytoplasmic (green shading) sources of invagination resistance. (B, C) Length of the furrow over time under a constant applied stress. The elastic and viscous coefficients were chosen arbitrarily to demonstrate characteristic behaviors. The initial and final *V*_*final*_ velocities are included for visualization purposes (dash-dotted black lines). In B, the total furrow length is the sum of the internal Burger body lengths (Eqs. 13, 14, 15). The colors of the plotted lines correspond with the boxed elements in panel A. In C, the furrow velocity reduces when limited by cytoplasmic drag (Eq. 18, highlighted green in A).

Next, we consider how the system behaves when it is submerged in a fluid, such that drag plays a role (Fig. 2A, C). Under constant stress, drag drives the furrow velocity to decay as Eq. 18, 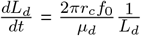. By noting that *L*_*d*_ = *L*_*T*_ – i.e., the drag force is parallel to the scaffold force (Fig. 2A) – we can conclude that our system will always transition from a fast to a slow velocity due to both the Burger body and the drag, which is the opposite of the slow-to-fast transition found in experiments. The conclusion of this result holds for constant, positive material coefficients. Therefore, constant material properties cannot capture experimental observations and a slow-to-fast transition requires dynamic changes to the material properties.

### 4.3 A dynamic scaffold produces the slow-to-fast shift

Next, we identify which parameters may be changed over time to achieve the slow-to-fast shift. We begin by considering cytoplasmic viscosity (*µ*_*d*_), which represents the drag force from a viscous medium (*f*_*d*_ in Eq. 18). Note that the drag force is proportional to the furrow length (defined as the length of PM spanning from the apical-lateral region to the basal tip in Fig. 1B, E) and velocity. Hence, we could associate a decrease of *µ*_*d*_ with an increase in furrow velocity and thus achieve a slow-to-fast shift. We assume that, as the furrow-bound scaffold invaginates through a viscous medium, it generates friction with the cytosol and other molecules that interact with this scaffold, such as microtubules. Thus, a decrease in scaffold density or cytosolic viscosity per unit length would decrease the drag force by decreasing *µ*_*d*_. However, to the best of our knowledge neither the density of the scaffold on the furrow nor the viscosity of the cytosol decrease with time. For these reasons, we hypothesize that the major contributor to the softening of material properties must be due to the scaffolding and not drag. Note, the net motor protein force is not tested as a possible mechanism of the slow-to-fast velocity shift because empirical evidence has shown that the scaffold softens concomitantly with the slow-to-fast shift, and is, therefore, a more justified mechanism to investigate [13].

Figure S1D visually demonstrates how fast-phase scaffold softening may be achieved using a Burger body (see Section S1.1). Panels D2 and D3 show that our options are limited to either (1) fast-phase softening (orange dashed lines), or (2) temporary slow-phase stiffening (orange dotted lines) to produce a slow-to-fast transition. We can constrain our options, however, by implementing the finding that viscosity *ζ* scales as a function of elasticity *R* as in equation 2 [13], given by

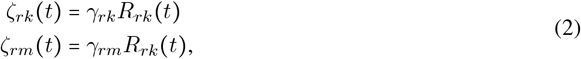

where *γ* is a scaling constant. Therefore, the Kelvin-Voigt body and Maxwell dashpot must change together.

We directly tested our hypothesis that a softening scaffold can cause the slow-to-fast shift by setting the elastic stiffness (*R*_*rk*_) in the fast phase to be less than in the slow phase (Fig. 3). The slow-phase KV elastic coefficient, *R*_*rk*0_, was set equal to the value measured by D’Angelo *et al*. [13], while the viscous coefficients in the Burger body, *ζ*_*rk*_ and *ζ*_*rm*_, were set with the elastic-viscous relation defined in equation 2 [13]. Whereas D’Angelo *et al*. [13] discovered the elastic-viscous correlation, we propose that viscosity is causally governed by the elastic elements, and not the reverse [22]. With defined material properties, the basal net motor protein force, *f*_*nmp*0_, remained our one free parameter, chosen such that the slow phase furrow invagination velocity equaled the value found in Figard *et al*. [3]. The value of *f*_*nmp*0_ remained constant for all of the remaining simulations (Table S1). Rather than use the fast phase KV elastic coefficient found by D’Angelo *et al*. [13], we chose *R*_*rk*_ to match the fast phase velocity [3] (Eq. 32). We solved for the fast phase *R*_*rk*_ this way because furrow invagination velocity is a direct measurement and therefore we judged it as more accurate than the measurement of material properties, which is influenced by the initial assumption of the type of viscoelastic body [3, 13]. Finally, we assumed that there was sufficient wrinkled membrane to produce a full length furrow; this allowed us to segregate the effects of the scaffold on furrow dynamics from the effects of the membrane.

**Figure 3:**
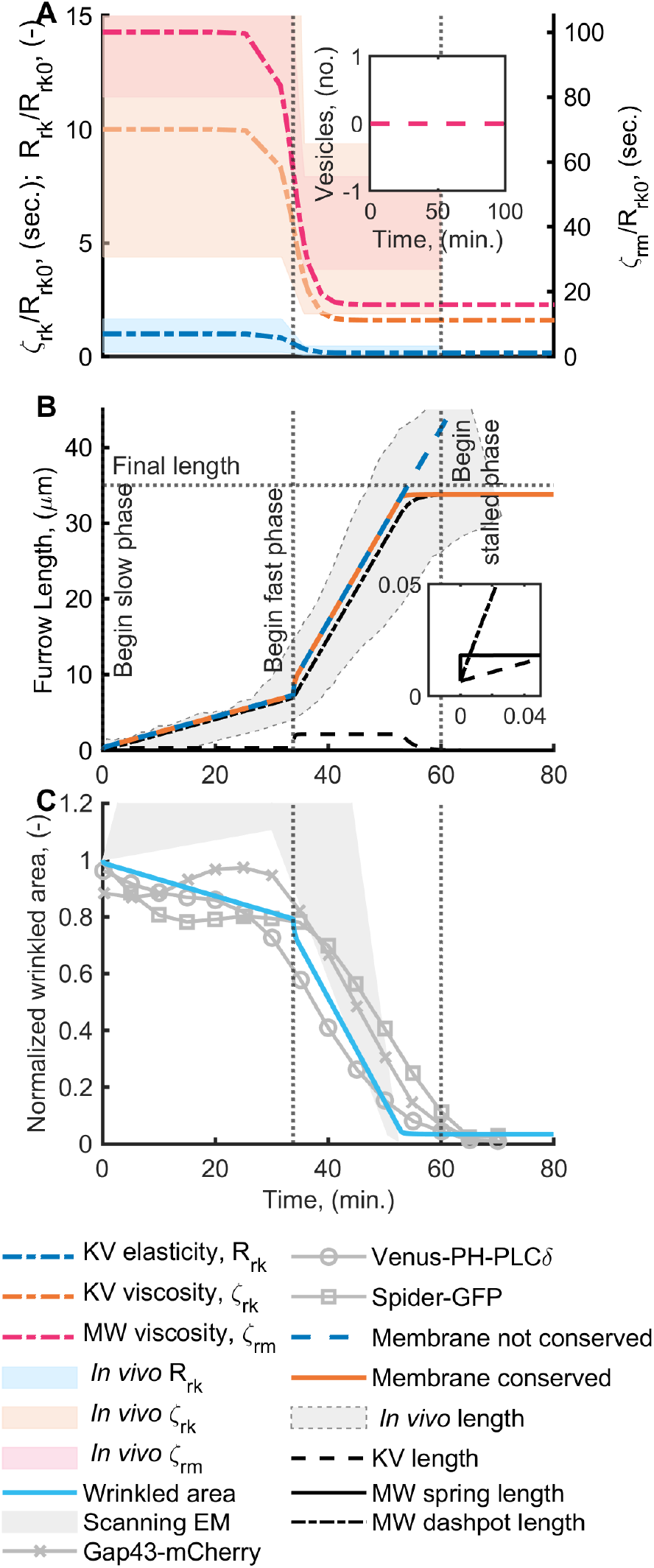
A viscosity-dominated scaffold with conservation of membrane area replicates furrow invagination kinetics. Figure 3. : (A) Time evolution of viscoelastic (VE) coefficients, from simulation (lines) and *in vivo* data (shaded) *ζ*_*rk*_, *ζ*_*rm*_, and *R*_*rk*_ from D’Angelo *et al*. [13]. Inset: (input) viscoelastic effects were isolated by initiating the wrinkled membrane reservoir with enough membrane to produce a 35 µm furrow. Therefore, no vesicles were exocytosed. (B) Furrow length, time evolution with constant applied force and changing VE properties from (A). Note two cases: the membrane is conserved (orange solid line) and not conserved (blue dashed line). *In vivo* length (shaded), *Final length, Begin fast phase*, and *Begin stalled phase* are from Figard *et al*. [3]. Inset: zoom showing the small displacement of the Maxwell spring (solid line); the dashdotted and dashed lines are the Maxwell dashpot and the Kelvin-Voigt body, respectively. (C) Wrinkled area over time, normalized by the value at *t* = 0 minutes. Experimental data of wrinkled membrane area are from measurements of the microvillus reservoir on the apical surface of the plasma membrane. Scanning electron microscopy data is approximated in Section S1.3 and raw data is from Figard *et al*. [3] (grey shaded). Fluorescence data, Gap43-mCherry, Venus-PH-PLC*δ*, and Spider-GFP are from Figard *et al*. [3] (gray lines). The blue line corresponds to the model. All other parameters used in the simulation are listed in Table S1. The wrinkled membrane area, KV spring, and furrow length RMS errors are .150, .117, and .035, respectively.

Our simulation indicates that the rate of furrow invagination is dominated by the scaffold and that the furrow length is predominantly stored as visco-plastic (permanent) deformation (Maxwell dashpot, Figs. 1D, 3B). This finding is in agreement with *in vivo* viscoelastic measurements of the scaffold during cellularization [13, 14, 26]. In our model, scaffold viscosity was scaled as a function of the Kelvin-Voigt elasticity (Eq. 2). So, while the Maxwell dashpot was the dominant contributor of furrow length, the Kelvin-Voigt spring was the ultimate controlling mechanism on furrow velocity. Finally, the Maxwell spring contributes no meaningful effect on furrow velocity nor length (Fig. 3B). Critically, this result builds confidence in the accuracy of our model because our free parameter in the fast phase, *R*_*rk*_, agrees with experimental measurements of scaffold stiffness [13] (Figs. 3A and S6). We observe that while softening of the scaffold in the fast phase allows for the slow-to-fast transition, this version of the model (Fig. 3B, blue dashed line) does not stop the furrow at 35 µm. Finally, in Fig. 3C our simulation closely matches experimental data of the apical membrane area [6]

### 4.4. Membrane area limits furrow length and replicates the fast-to-stalled shift

Next, we consider how the plasma membrane may be used to stop furrow invagination at the fast-to-stalled transition (Fig. 1C). To justify wrinkled membrane area depletion as a stopping mechanism, we note that furrow length and microvillus membrane are tightly coupled – stalled furrow invagination halts microvilli depletion and microvilli depletion stalls furrow invagination [3, 6]. In Figure 3B, when membrane area is not conserved (dashed blue line), the furrow continues to lengthen because both the drag and the Burger body resistance forces are functions of velocity over long timescales; they behave as fluids, which do not have a defined stopping length. As mentioned previously, a fluid-like cytoskeleton is in agreement with experimental findings [13, 14]. Therefore, we concluded that the model in Fig. 3B (dashed blue line) was missing a stalling mechanism, and we sought guidance from experimental evidence.

At the end of cellularization, the apical surface transitions from a bumpy, microvillus surface, to a smooth dome (Fig. 1E) [3]. Furthermore, in experiments that limit the amount of apical microvilli, the final furrow length reduces proportionally with microvilli density [6]. Therefore, we use our model to test the hypothesis proposed by Figard *et al*. [6], that furrow length is limited by available membrane area. To implement this feature, we conserve membrane area by allowing it to exist either as a wrinkled area reservoir or unwrinkled membrane along the furrow length (Fig. 1E, Eqs. 20, 21, and 22, Section 6.3). Phenomenologically, we propose that as wrinkled area, *A*_*w*_, approaches 0 µm^2^, the membrane stress increases as

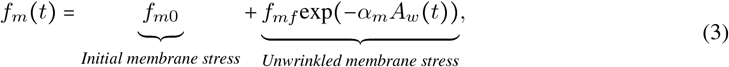

where *f*_*mf*_ and *α*_*m*_ are scaling constants. The membrane limitation on furrow length is added as a parallel force to the pre-existing resistive forces of the scaffold and cytoplasmic drag (Eq. 1, see the orange line in Fig. 3B for the corresponding membrane-limited length). The complete force balance is shown schematically in Fig. 1D. Our goodness-of-fit was measured with the root mean square error (RMSE) of the residuals between the normalized experimental mean and model data at 5-min. intervals (see Figs. S5, S6, and S7 and the caption in Fig. 3). A conservation of membrane area stops the furrow length at ∼ 35 µm due to a loss of wrinkled membrane. Our model supports *in vivo* findings that membrane can limit furrow length [6], and proposes that the fast-to-stalled length may be governed by a loss of wrinkled membrane area.

### 4.5. Scaffold assembly and disassembly replicate the slow-to-fast shift

Thus far we have modeled complete furrow invagination with a single set of equations (Fig. 3). Here, depletion of the wrinkled membrane reservoir caused the invagination to stall (the fast-to-stalled shift). Additionally, we found that softening of the scaffold alone is sufficient to replicate the switch from the slow to the fast velocity. This result is supported by measurements of scaffold softening at the slow-to-fast shift [13]. Moreover, it challenges the hypothesis that membrane scarcity governs invagination velocity [16]. However, it is not yet well understood what mesoscale mechanisms may cause the scaffold to soften.

In sections 4.5.1 and 4.5.2, we propose that the scaffold softens due to a change in the network organization of its constituent cytoskeletal components [14]. For example, consider the effect of network architecture when a tensile load is applied to a bundle of actin filaments as compared to a network of branched actin filaments. While both are constructed of identical filaments, the bundled network can sustain greater tensile loads than the branched network. We use our model to simulate the viscoelastic changes of the scaffold under simplified configurations: when cytoskeletal components are connected (1) in series (end-to-end), such that each filament is connected to its neighbor in a chain-like fashion, extending along the long axis and (2) in parallel, such that the filaments are stacked on top of each other with their ends aligned.

Additionally, we propose that the reorganization of the cytoskeletal network is closely linked with vesicle exocytosis to the PM. To rationalize this mechanism, the microvilli that support furrow invagination require both membrane and cytoskeletal components to form [3]. Furthermore, as additional microvilli are assembled it has been suspected that cytoskeleton or cytoskeletal regulators of the cytoskeleton are transported alongside vesicles to the PM during cellularization, although the precise mechanisms remain unclear [16, 18, 19]. Finally, we hypothesize that vesicle transport is implicated in the slow-to-fast invagination shift because scaffold stiffness drops when vesicle exocytosis stops – at the slow-to-fast shift [13]. In our model, the timing cytoskeletal element assembly into the scaffold (*n*_*mvc*_) can be simultaneous with vesicle exocytosis 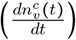,

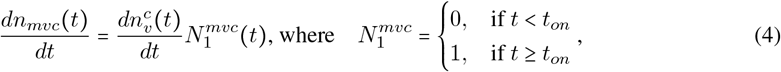

or with a delay such that new cytoskeletal elements are assembled between times *t*_*on*_ and *t*_*off*_,

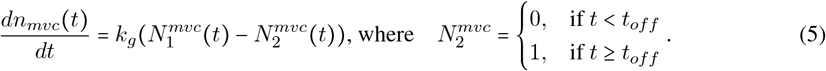

Here, cytoskeletal assembly begins and ends at times *t*_*on*_ and *t*_*off*_, respectively, and *k*_*g*_ is the time-independent form of, 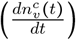. Importantly, equation 4 allows membrane addition to the PM without any associated cytoskeletal assembly whenever *t*_*on*_ is greater than the start time of exocytosis. We do not assume that cytoskeletal components are released by vesicles, only that their transport pathways are temporally coupled in order to assemble the PM-cytoskeleton composite. Equation 5 and an expanded form of equation 4 can be found in section 6.5.1 (Eqs. 31 and 30). Finally, the transport of wrinkled membrane and cytoskeleton to the PM is non-spatial because we are using a system of ODEs in time. Specifically, wrinkled membrane and cytoskeletal (scaffold) viscoelasticity are stored as the quantities, *A*_*w*_ (Eq. 22) and *R*_*rk*_ (Eqs. 6 and 7), respectively, that influence the furrow velocity through the governing equations (Eq. 36). For the remainder of this paper, we explore how viscoelasticity of the scaffold is affected by the dynamic coordination of exocytosed membrane and cytoskeletal assembly.

#### 4.5.1. Tension-induced scaffold disassembly replicates the slow-to-fast shift

In our first proposed mechanism, we softened the scaffold by allowing the tension from motor proteins to disassemble the cytoskeletal elements of the scaffold (See Fig. 4A-C for a schematic of the model. See Fig. S1 and section S1.1 for a mechanistic understanding of how the removal of parallel elements softens the scaffold). This choice is supported by evidence that the disassembly rate of molecules increases when tension is applied [14, 27–29]. Thus, the parallel elements are disassembled as shown in Figure 4B (see also Supplementary Material S1.1, Fig. S1A-C). The total elastic scaffold stiffness *R*_*rk*_, given by

**Figure 4:**
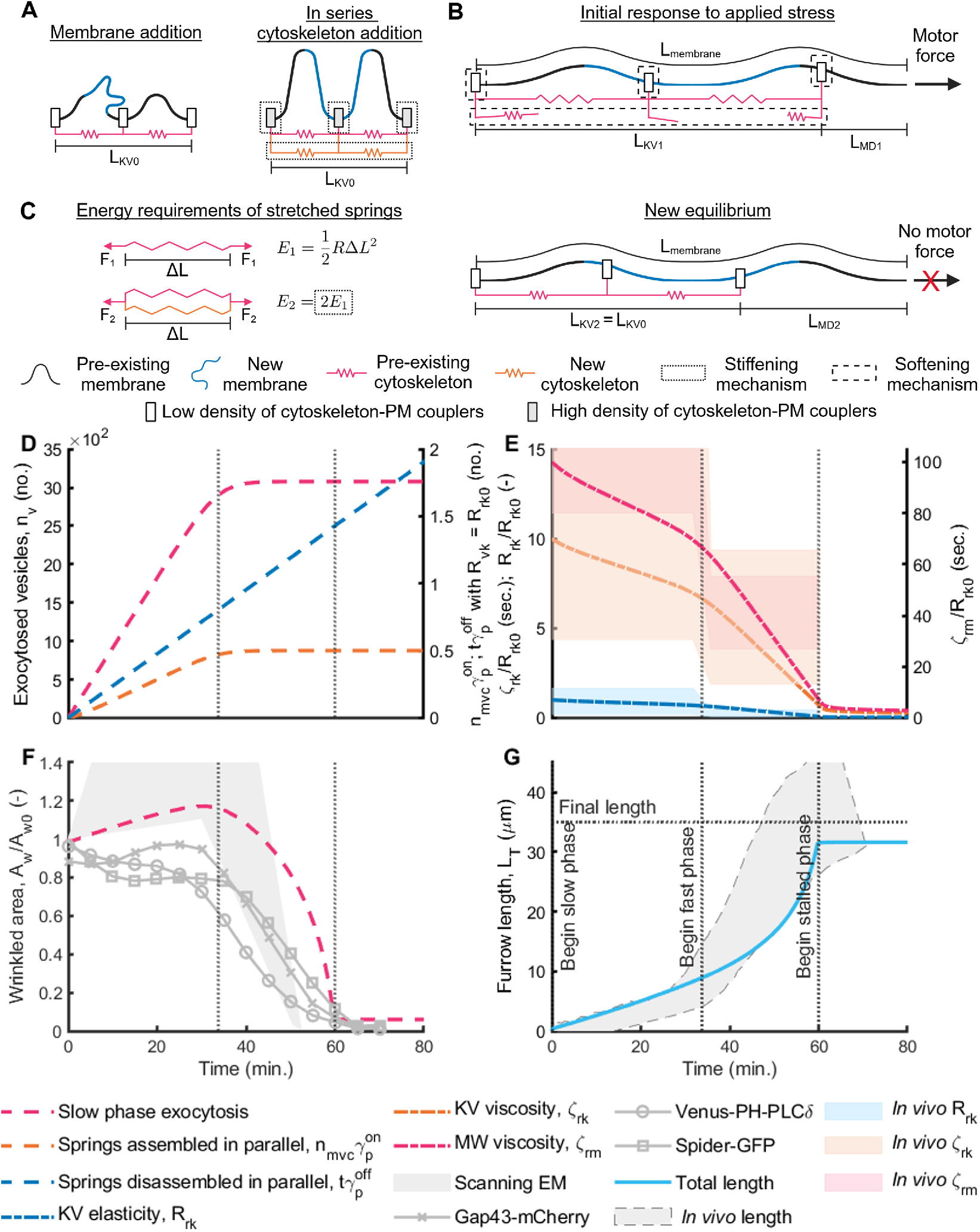
Vesicle-coupled scaffold assembly paired with tension-coupled disassembly replicates *in vivo* results. Figure 4: A schematic of wrinkled membrane and scaffold assembly (A) and disassembly (B). (C) Mathematical proof of energy requirement changes due to spring assemble. Two springs assembled in parallel (bottom), require two times as much energy as a single spring when stretched equal lengths, Δ*L*. Compare the energy values of one spring, *E*_1_, to two springs, *E*_2_. *F*_1_ and *F*_2_ are the corresponding forces, where Energy = Force x Distance. (D) Vesicle exocytosis (dashed magenta line) and the number of parallel springs assembled (dashed orange line) and disassembled (dashed blue line) over time. Exocytosis was simulated with equation 24, *N*_*off*_ = 1 (unitless) for all time, Δ*t*_*g*_ 33. = 75 minutes, and *L*_*re*0_ = 0 µm. The number of assembled and disassembled springs were simulated with 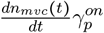 and 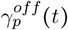, respectively, and found in the first and second terms of equation 35a, where 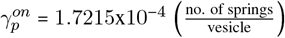 and 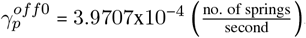. The coupling between cytoskeletal assembly and vesicle exocytosis was simulated with equation 30, where *t*_*on*_ 0 minutes (Table S2). (E) The time evolution of viscoelastic properties. These properties were governed by equation 6. Shaded areas correspond to the *in vivo* viscoelastic coefficient data, *ζ*_*rk*_, *ζ*_*rm*_, and *R*_*rk*_ [13]. Model data is represented with dash-dotted lines. (F) Wrinkled area over time, normalized to its initial value. The dashed magenta lines corresponds to slow phase exocytosis, as in panel D. Scanning EM data is approximated in Section S1.3 and raw data is from Figard *et al*. [3] (gray shaded area). Fluorescence data, Gap43-mCherry, Venus-PH-PLC*δ*, and Spider-GFP are from Figard *et al*. [3] (gray lines). (G) Furrow length over time. The gap between the *Final length* (horizontal dotted lines) and model *Total length* (light blue line) at the end of cellularization is a numerical artifact. Length data, *In vivo* length, *Final length, Begin fast phase*, and *Begin stalled phase* are from Figard *et al*. [3]. Viscoelastic properties begin changing at, *t*_*on*_ = 0 minutes. All other parameters used in the simulation are listed in Table S1. The apical membrane area, KV spring, and furrow length RMS errors are .357, .189, and .151, respectively (see Figs. S5, S6, and S7 and the caption in Fig. 5).

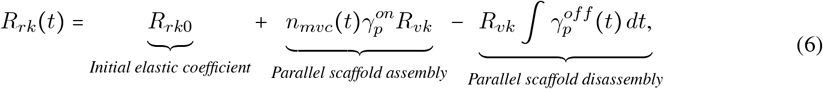

was softened with the third term by integrating the spring disassembly rate, 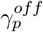, over time. Here, the integral is left unevaluated because 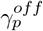 is later redefined to simulate *in vivo* experiments that varied the tension (and therefore the spring disassembly rate) felt by the scaffold (relevant definitions can be found in Eqs. 35, and 9 - 11). Without an experimentally measured stiffness of individual cytoskeletal springs, *R*_*vk*_, we set each disassembled spring equal to the initial stiffness of the total scaffold, *R*_*rk*0_, measured by D’Angelo *et al*. [13]. Therefore, this choice implies that each disassembled spring represents a fraction of the total initial network of springs in the scaffold. The third term also implies that scaffold disassembly reduces the density of cytoskeletal elements (parallel disassembly). Biologically, this can be understood as unbinding crosslinking and transmembrane proteins from cytoskeletal filaments like F-actin (Fig. 4B). Our model homogenizes the effects of these crosslinkers and transmembrane proteins because that is the limit of resolution of measured scaffold material properties (tens of µm) [13, 14].

The second term of equation 6 allows the scaffold to stiffen by the addition of parallel PM-bound cytoskeletal elements (Fig. 4A). Here, 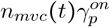 is the total number of new springs assembled in parallel, with an elastic coefficient of *R*_vk_ = *R*_*rk*0_, as was done in the third term.

Substituting the disassembly equation (Eq. 6) into the governing equations (Eqs. 36) of the force balance (Eq. 1), we found that parallel scaffold assembly and disassembly rates competed to produce the slow-to-fast shift in invagination velocity (Figure 4D-G). In Fig. 4D, vesicles were exocytosed to the PM for the first 33.75 minutes of cellularization with equation 24 (dashed magenta line) [6, 17]. Parallel springs were assembled simultaneously with vesicle exocytosis for the duration of exocytosis (Fig. 4A and C, Eq. 5, *t*_*on*_ = 0 minutes). We tracked the contributions of scaffold assembly 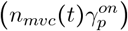 and disassembly 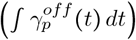 terms by independently plotting the second (dashed orange line) and third (dashed blue line) terms from equation 6, respectively (Fig. 4D). Our only free parameters were the number of springs assembled per vesicle exocytosed, 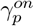, and the spring disassembly rate, 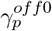 (Eq. 35). We evaluated the quality of the disassembly-softening mechanism by comparing the model to experimental measurements of apical viscoelasticity (Fig. 4E), apical membrane area (Fig. 4F) and furrow length (Fig. 4G) as functions of time. We found that disassembly alone could not produce the slow-to-fast shift (Fig. 4E and G). Rather, it was necessary for the scaffold to also stop assembling stiffening elements. This finding is made most clear in Figure S2, which plots furrow length as a function of time, when assembly and disassembly rates are varied. Finally, as shown in Figure 4F, wrinkled area accumulation in the slow phase agrees with scanning electron microscopy data, but predicts greater apical membrane than fluorescent data [3, 6].

#### 4.5.2. Scaffold assembly replicates the slow-to-fast shift in furrow invagination velocity

In our second proposed mechanism, we softened the scaffold by assembling cytoskeletal elements in series during the fast phase of cellularization (See Fig. 5A, B for a schematic of the model. See Figs. 5C and S1 and section S1.1 for a mechanistic understanding of how series addition softens the scaffold). This choice is supported by the well-studied formation of new microvilli during cellularization, where the cytoskeleton is needed to stabilize the wrinkled shape of the membrane [3, 6, 17]. As mentioned above, it is suspected that cytoskeleton or cytoskeletal regulators are transported alongside vesicles to the PM during cellularization, although the mechanisms remain unclear [16, 18, 19]. To capture these effects, the elastic scaffold stiffness, *R*_*rk*_,

**Figure 5:**
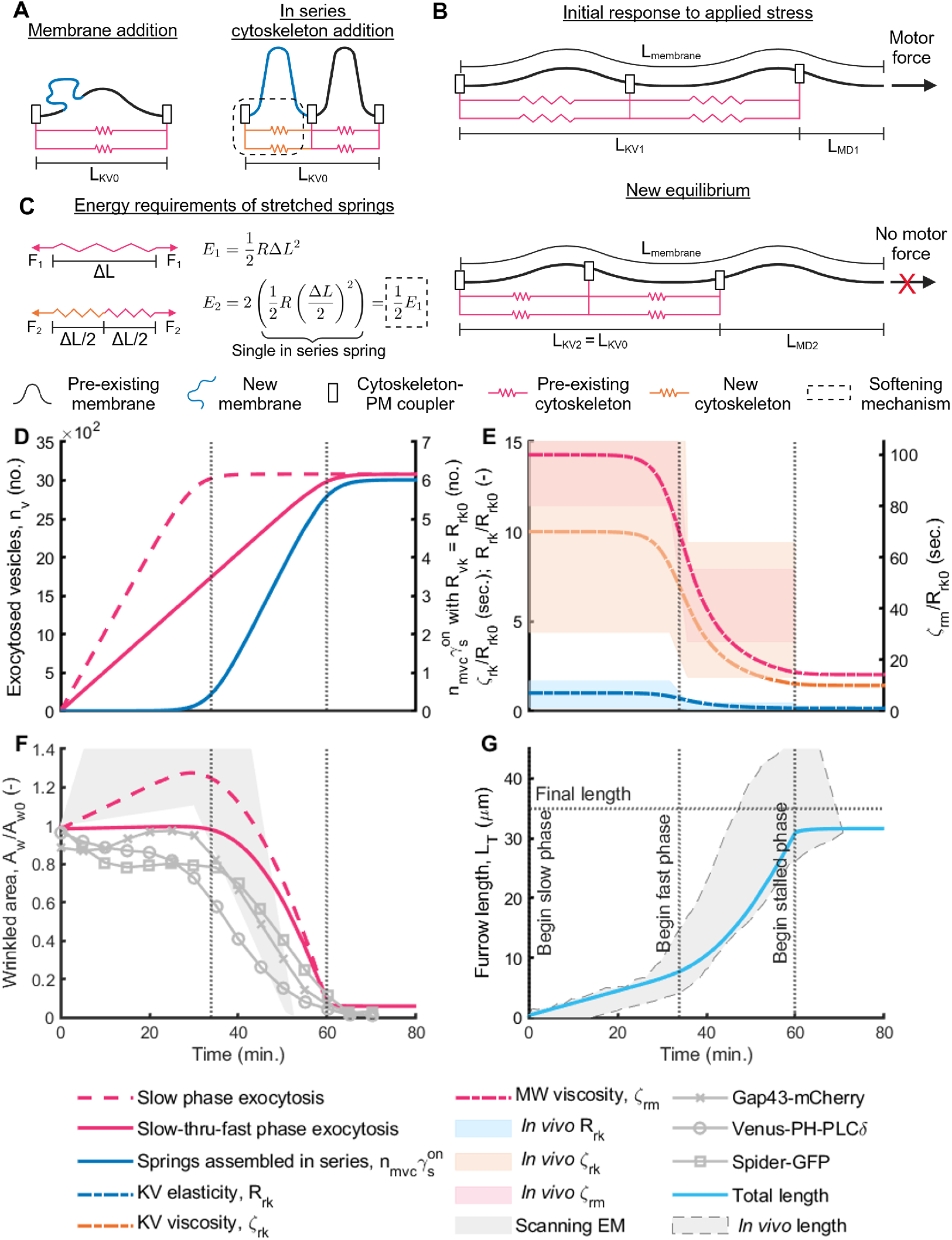
Vesicle-coupled scaffold assembly replicates *in vivo* results. Figure 5. : A schematic of wrinkled membrane and scaffold assembly (A) and furrow lengthening (B). (C) Mathematical proof of energy requirement changes due to spring assemble. Two springs assembled in series (bottom), require half as much energy as a single spring when stretched equal lengths, Δ*L*. Compare the energy values of one spring, *E*_1_, to two springs, *E*_2_. *F*_1_ and *F*_2_ are the corresponding forces, where Energy = Force x Distance. (D) We consider two cases for vesicle exocytosis over time: in the slow phase (dashed magenta line, Δ*t* = 33.75_*g*_ minutes) and through the slow and fast phases (solid magenta line, Δ*t*_*g*_ = 60 minutes). However, both mechanisms lead to the same rate of in series spring assembly (blue solid line). To do this, the timing of spring assembly relative to vesicle exocytosis was different. The rate of cytoskeletal assembly was governed by Eqs. 31 and 33 for *slow phase exocytosis*, where 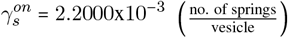. For *slow-thru-fast phase exocytosis*, Eqs. 30 and 33 are used, where 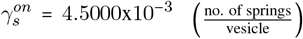 (dash-dot lines represent both cases). Both “Slow phase exocytosis” and “Slow-thru-fast phase exocytosis” added membrane to the PM using Eq. 24, with *N*_*off*_ 1 (unitless) for all time and *L*_*re*0_ 0 µm. (E) The time evolution of viscoelastic properties. These properties were governed by equation 7. Viscoelastic properties begin changing at, *t*_*on*_ 33.75 minutes. The shaded areas correspond to the *in vivo* viscoelastic coefficient data, *ζ*_*rk*_, *ζ*_*rm*_, and *R*_*rk*_ [13]. (F) Wrinkled area over time. The magenta lines correspond to the slow and slow-thru-fast phase exocytosis cases as in panel (D). Scanning electron microscopy data is approximated in Section S1.3 and raw data is from Figard *et al*. [3] (gray shaded area). Fluorescence data, Gap43-mCherry, Venus-PH-PLC*δ*, and Spider-GFP are from Figard *et al*. [3] (gray lines). (G) Temporal evolution of furrow length. The model (blue line) is from the “slow-thru-fast exocytosis” case, although the “slow phase exocytosis” produces an equivalent curve. The gap between the *Final length* (horizontal dotted lines) and the model *Total length* at the end of cellularization is a numerical artifact. Length data, *In vivo* length, *Final length, Begin fast phase*, and *Begin stalled phase* are from Figard *et al*. [3]. All other parameters used in the simulation are listed in Table S1. For the *slow phase exocytosis* model, the apical membrane area, KV spring, and furrow length RMS errors are .368, .138, and .127, respectively. For the *slow-thru-fast phase exocytosis* model, the apical membrane area, KV spring, and furrow length RMS errors are .242, .135, and .129, respectively (see Figs. S5, S6, and S7 and the caption in Fig. 5).

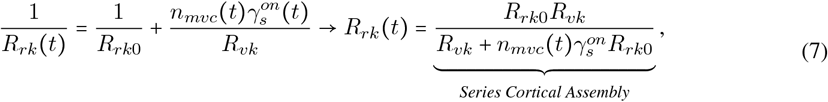

was softened as the number of assembled cytoskeletal elements, 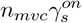, increased. Where each new spring assembled in series had an elastic coefficient of *R*_*vk*_ = *R*_*rk*0_, as in section 4.5.1. Our only remaining free parameter, 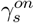, is tuned to capture the effective change in stiffness per vesicle.

Importantly, to produce the slow-to-fast shift, scaffold softening with equation 7 must only occur in the fast-phase (Fig. 5D, blue line). To achieve fast phase softening, we simulated assembling springs in series both simultaneously (Eq. 4) and delayed (Eq. 5) with respect to vesicle addition. First, vesicles were exocytosed only in the slow phase with cytoskeletal assembly delayed by 33.75 minutes relative to membrane addition (Eq. 5, Fig. 5D, dashed magenta line). Then we repeated the simulation, but vesicles exocytosed for the duration of cellularization (Fig. 5D, solid magenta line). Here, only vesicles were exocytosed to the PM for the first 33.75 minutes, afterwhich both vesicles and cytoskeletal elements were assembled into the composite PM-scaffold. Evidence in support of exocytosis only in the slow-phase came from experiments that showed furrows reached full length with only slow-phase Golgi-derived vesicle exocytosis [6]. However, evidence in support of exocytosis for the duration of cellularization came from experiments that showed vesicle exocytosis into the apical-lateral region of the furrow during the fast-phase [2, 17]. In summary, Figure 5 aims to test first, if (1) scaffold assembly is a plausible softening mechanism, and (2) for what duration vesicles should be exocytosed.

Substituting Eq. 7 into the governing equations (Eqs. 36) of the force balance (Eq. 1), we find that in series scaffold assembly can produce the slow-to-fast shift in invagination velocity (Figure 5D-G). This slow-to-fast shift is produced with slow and slow-thru-fast phase exocytosis. We obtained the same number of springs assembled in series in both cases by changing the parameter 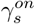. We evaluated the quality of the assembly-softening mechanism by comparing the model to experimental measurements of apical viscoelasticity (Fig. 5E), apical membrane area (Fig. 5F) and furrow length (Fig. 5G) as functions of time. We found that exocytosis of vesicles for the duration of cellularization better represented fluorescent measurements of wrinkled area accumulation (Fig. 5F, solid magenta line), while, “slow phase exocytosis” better represented scanning electron microscopy wrinkled area accumulation (Fig. 5F, dashed magenta line) [3, 6].

### 4.6. The slow-to-fast shift is driven by the coordinated transport of both vesicles and new cytoskeletal elements to the PM

Thus far, we have identified plausible mechanisms of scaffold softening by way of cytoskeletal disassembly and assembly (Figs. 4 and 5). We evaluated each of these mechanisms against furrow length, apical area, and viscoelastic coefficients. All of our model comparisons were focused on experiments that measured features of furrow invagination under normal cellularization conditions. We next investigated if the models defined in Figures 4 and 5 could capture experimental observations when normal cellularization conditions were disrupted. First, in section 4.6.1 we tested the role of the assembly and disassembly softening mechanisms when motor protein forces are increased [13]. Then, in section 4.6.2 we tested the model when the timing of vesicle transport was disrupted [6].

#### 4.6.1. The slow-to-fast shift is dominated by scaffold assembly and not disassembly

In this section, we quantified the degree of scaffold softening due to an increased applied motor force. By comparing the fold-change of simulated scaffold stiffness to *in vivo* data [13], we scrutinized the predictive power of our scaffold softening mechanisms (Figs. 4 and 5). Here, we sought to replicate the following *in vivo* experiment: researchers attached a ferrous bead to the apical surface of an invaginating furrow and then pulled the bead parallel to the apical surface with a magnet (Fig. 6A) [13]. This test showed that the apical viscoelastic moduli were independent of both the frequency and amplitude of an applied force. To simulate the experimental input with our system of ODEs, we assumed that the force exerted by the bead increased the tension on the scaffold. Therefore, in our model, we increased the applied motor protein force (Eqs. 10 and 11) with frequencies and amplitudes equal to *in vivo* values (Fig. 6B). Figure 6C shows the four different force curves used to simulate *in vivo* inputs. First, our control case used the same *f*_*nmp*_ value as all previous simulations (‘Constant, low’, Eq. 9 implies Δ*f* = 0). Next, we increased the amplitude of Δ*f* to match the smallest Δ*f* input by D’Angelo *et al*. [13] (‘Constant, high’, Eq. 11). Then, we oscillated *f*_*nmp*_ between our ‘constant, low’ and ‘constant, high’ cases to evaluate the effects of frequency on scaffold softening (‘Oscillatory’, Eq. 10). Finally, we increased Δ*f* to within one order of magnitude of our ‘constant, high’ case to examine the sensitivity of the model to moderate force variation (‘Constant, order of mag.’, Eq. 11). All other parameters and equations matched those used in Figures 4 and 5. The first outputs of our model were the normalized elastic stiffness, *R*_*rk*_ /*R*_*rk*0_, for the assembly (Fig. 6D) and disassembly (Fig. 6E) softening mechanisms, where *R*_*rk*0_ is the initial stiffness of the scaffold. Note that the viscosity is linearly coupled with the KV elastic stiffness (Eq. 2), and hence, we did not include this plot. Our second output calculated the fold-change of elasticity at each point in time (Fig. 6F). To do this, we divided our test cases (‘constant, high’, ‘oscillatory’, and ‘constant, order of mag.’) by our ‘constant, low’ control case, 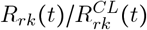 Here, 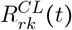 is the elastic coefficient for the ‘constant, low’ case. We emphasized calculating the fold-change at each point in time by including (*t*). This method exposed the difference in the expected verses the actual elastic stiffness under different motor protein force inputs.

**Figure 6:**
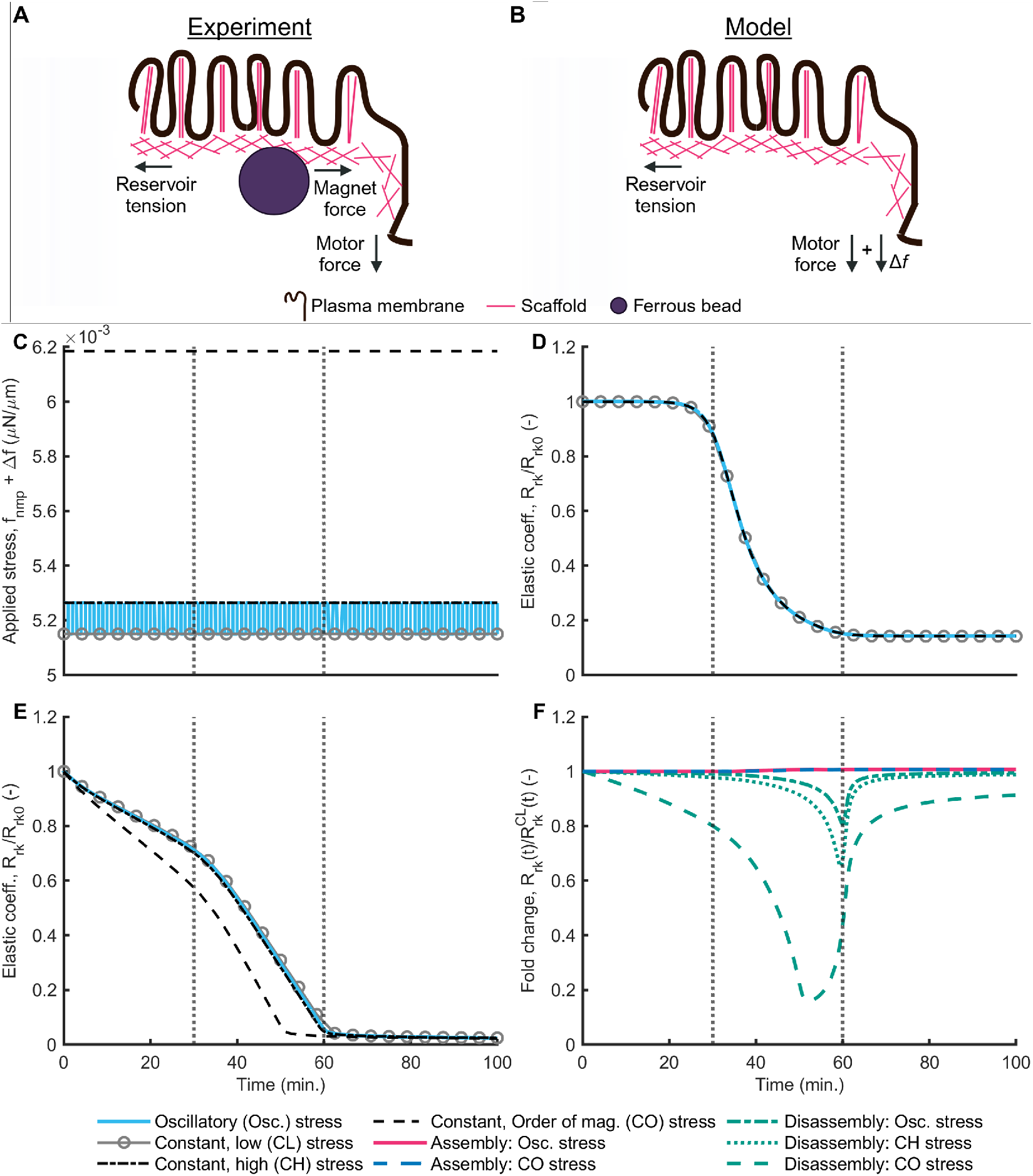
An assembly-, and not disassembly-softened scaffold, is required to match the experimental invariance to force amplitude and oscillation. Figure 6. : A schematic representation of the forces felt by the scaffold in the experiment (A) and how they translate to our model (B). (C) Temporal evolution of the forces applied to the system at a constant low (control, CL) (Eq. 9), a constant high (CH) (Eq. 11), an oscillatory square-wave (Osc.) (Eq. 10), and a constant, same order-of-magnitude (CO) amplitude (Eq. 11). The increased force, *δ*_*f*_, of the CH and Osc. tests matched D’Angelo *et al*.’s [13] maximum applied stress (Table S1). Elastic coefficient evolution over time for the scaffold assembly (D) and disassembly (E) softening cases. The model and parameter values for panels D and E were identical to Figs. 5 and 4, respectively. All other parameters used in the simulation are listed in Table S1 and Table S2. (F) Fold-change of the elastic coefficient, evaluated at each time-point of the Osc., CH, and CO stress cases relative to the CL case.

In Figure 6D, the viscoelastic behavior of the assembly-softened scaffold (Fig. 5) demonstrated that this mechanism is invariant to tension amplitude and frequency. This is consistent with the observations *in vivo*[13] (Fig. 6D, F). This force invariance is because softening was driven by springs assembled in series as outlined in Figures 5A, C and S1A, B – which makes this softening mechanism entirely insensitive to any applied force (Eq. 7).

In comparison, we observed that the disassembly-softened scaffold in Figure 6E, is highly dependent on the *in vivo* tensile load (Fig. 6E, F). By design, the disassembly mechanism softens as a function of force, so some amount of softening was expected. Importantly, we were interested in discovering whether a physiologically relevant change in elasticity would occur under experimental loading conditions. At the low end of the magnetic force (Δ*f*), scaffold stiffness experienced a fold-change reduction between 20% and 30% (Fig. 6F). Furthermore, within one order of magnitude of the low end of the magnetic force, scaffold stiffness experienced a fold-change reduction of 80% (Fig. 6F). Altogether, this suggests that disassembly is unlikely to be a dominant mechanism of scaffold softening.

#### 4.6.2. A sub-apical reservoir governs the slow-to-fast shift by redirecting vesicles from the apical to the apical-lateral surface

Next, we investigated how blocked vesicle transport to the PM affects furrow invagination behavior. These simulations allowed us to scrutinize the types of coupling between cytoskeleton and membrane addition by comparing invagination length and velocity to *in vivo* data. As mentioned previously, our model does not assume the cytoskeletal components are transported on or in vesicles; it only assumes that there is temporal alignment in their transport [6, 16, 18–20]. Furthermore, because our model couples cytoskeleton assembly to vesicle exocytosis, blocking the latter will inhibit the former. Here, we sought to replicate the following *in vivo* experiment: researchers blocked Golgi export of vesicles by injecting the drug Brefeldin-A (BFA) in the embryo at the start of the slow and fast phases of cellularization (Fig. 7A1) [6]. To simulate this experiment with our system of ODEs, *t*_*bfa*_ in equation 25 represented BFA injections at 0 and 33.75 minutes. These injections had the effect of blocking cytoskeletal assembly due to the upstream inhibition of Golgi-dervied vesicles, 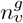. In Figure 7B and C, all other parameters and equations matched those used in the *slow phase exocytosis* and *slow-thru-fast phase exocytosis* cases of Figure 5, respectively. We did not simulate the disassembly softening mechanism in this section because it was found to be an unlikely scaffold softening mechanism in Fig. 6. Finally, vesicles were transported as shown in Figure 7Ai. Our model output reported furrow length as a function of time. We compared our BFA simulation cases to invagination velocities from Figure 5 (shown as the ‘no BFA’ case in Fig. 7) and stopping lengths to data from Figard *et al*. [6] (shown as blue and light blue dots in Fig. 7).

**Figure 7:**
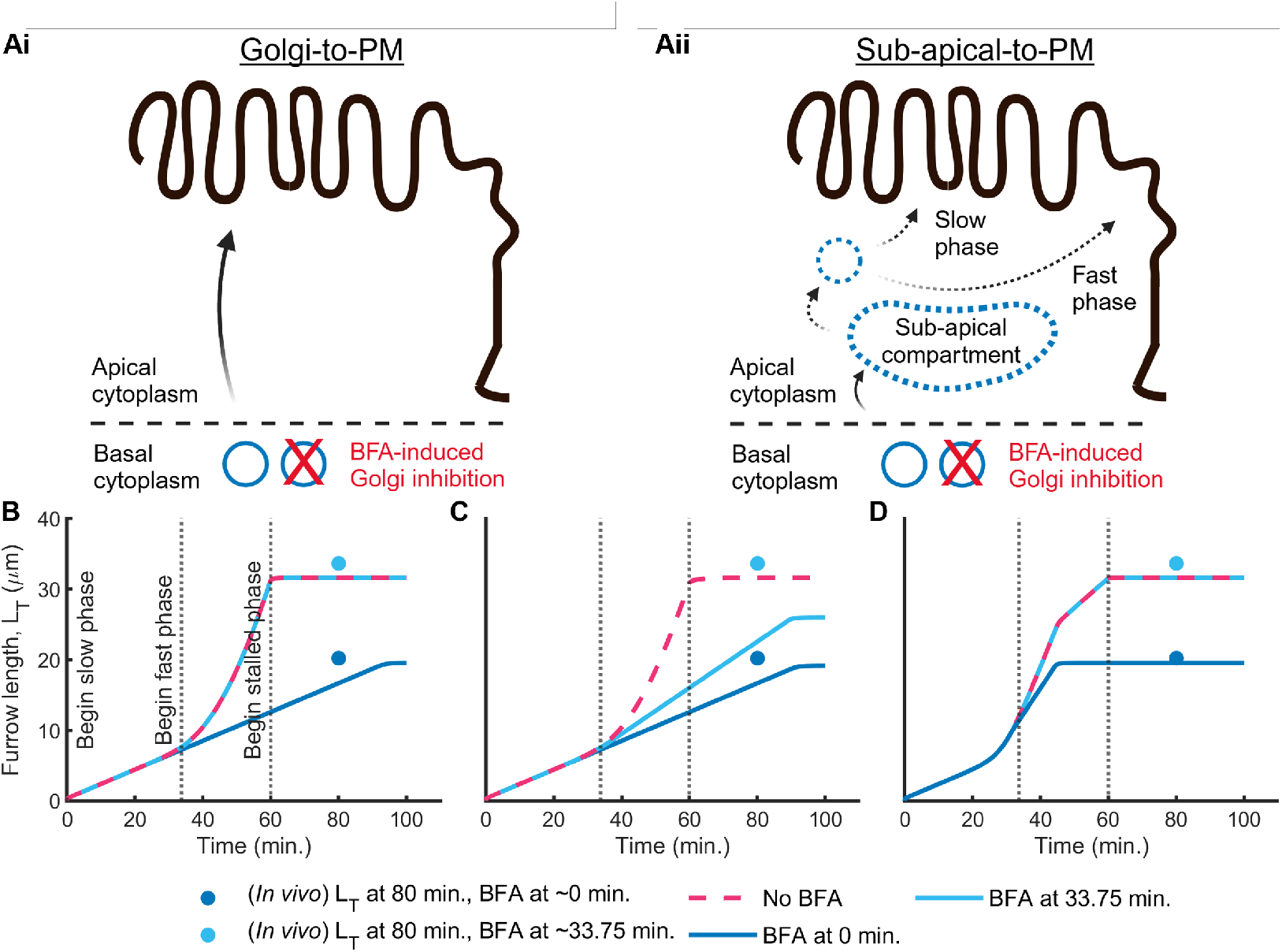
Golgi export through a sub-apical reservoir replicates Brefeldin-A (BFA) results. (Ai, Aii) A schematic representation of Golgi-derived vesicles originating in the basal cytoplasm of the embryo and enter the space of the apical cytoplasm. BFA inhibits the formation of Golgi-derived vesicles [6]. (Ai) Golgiderived vesicles are exocytosed directly to the PM. Panels B and C used this pathway. (Aii) Golgi-derived vesicles are first transported to a sub-apical compartment and then sub-apical vesicles were exocytosed to the PM. Panel D used this pathway. (B-D) The time evolution of furrow length, when Golgi-derived vesicle transport was stopped at *t*_*bfa*_ = 0 and 33.75 minutes into cellularization (Eq. 25). *In vivo* data is from Figard *et al*. [6]. Simulation in (B) extended the results from Figure 5, *slow phase exocytosis*, in which all vesicles were exocytosed during the slow phase (first 33.75 min.) and viscoelastic changes to the scaffold were delayed by 30 minutes. Excluding *t*_*bfa*_, all other parameters and equations are unchanged. Simulation in (C) extended the results from Figure 5, *slow-thru-fast phase exocytosis*, in which vesicles were exocytosed throughout cellularization (60 minutes) and viscoelastic changes to the scaffold take effect only in the fast phase (last 26.25 min.). Excluding *t*_*bfa*_, all other parameters and equations are unchanged. In (D), membrane was transported from the Golgi, through the sub-apical compartment, and into the PM domain with equations 24-29. The duration of Golgi export was Δ*t* = 33.75 minutes, *L* = 6_*g re*0_ µm, and the duration of sub-apical-to-PM exocytosis was Δ*t* = 45_*re*_ minutes. The membrane-viscoelastic coupling was governed by equations 30 and 34, where *t*_*on*_ = 33.75 min., and 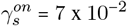. All other parameters used in the simulation are listed in Table S1.

The *slow phase exocytosis* case in Figure 7B was unable recover a slow-to-fast shift after BFA injections. Specifically, the 0-minute simulation failed to achieve the slow-to-fast transition as was found *in vivo* [6]. This is because scaffold softening is coupled to vesicle transport in the model, and therefore, with vesicle transport blocked at minute zero, no cytoskeletal elements were assembled in-series to soften the scaffold. However, in agreement with Figard *et al*. [6] the simulation did eventually reach a stopping length of 20 µm. The simulation stops short of the normal 35 µm furrow length because without vesicles to resupply the wrinkled membrane reservoir, the furrow length is limited to the initial supply of wrinkled membrane area. In the 33.75-minute BFA injection case, all vesicles were transported to the PM and so the invagination velocities and stopping length match the ‘no BFA’ case and *in vivo* data [6].

The *slow-thru-fast phase exocytosis* case shown in Figure 7C was unable to reproduce slow- and fast- phase velocities for the same reason as the *slow phase exocytosis* case. In addition to the 0-minute BFA case, here the 33.75-minute case also struggled to produce a meaningful slow-to-fast shift. From this, we can conclude that cytoskeletal transport in the fast phase is necessary to produce the slow-to-fast phase shift. This is true within the framework that the transport of vesicles and cytoskeletal elements are coupled by equations 4 or 5.

These failures indicate that the cytoskeleton-vesicle couplings defined in the assembly softening scaffold (Figure 5) are insufficient to replicate the results from Figard *et al*.’s [6] BFA study. To address these failures, we propose that an internal sub-apical compartment may function as a secondary reservoir to the external wrinkled membrane reservoir. Specifically, this sub-apical compartment could serve as an intermediate transit hub between the Golgi and the PM (Fig. 7Aii). In this scenario, for the 0-minute BFA injection, initial membrane stored in the sub-apical compartment would traffic to and add to the wrinkled membrane region of the PM to trigger the slow-to-fast shift. This could work because BFA targets Golgi-derived vesicles, thus leaving the sub-apical compartment unaffected. Then, for the 33.75-minute BFA injection, the slow-to-fast shift would be unaffected because the Golgi has been found to export all necessary membrane during the slow phase [6, 30]. Indeed, the Golgi-to-sub-apical pathway is supported by *in vivo* findings that the Golgi-derived transmembrane protein, Neurotactin, accummulates in the sub-apically positioned recycling endosome when vesicle budding is inhibited [21]. Also, there is a shift in vesicle exocytosis from the microvillus domain in the slow phase to the apical-lateral domain of the furrow in the fast phase [30]. Concomitant with the shift in exocytosis, Rab8 decorated vesicles that are destined for addition at the apical-lateral furrow domain become increasingly associated with sub-apical recycling endosome in the fast phase [2, 30]. As a final point, the apical-lateral domain continues to lengthen in the fast phase, despite the apical microvilli no longer being pulled into the furrow (see Lecuit & Wieschaus’s Fig. 4C [17]). Therefore, if the furrow membrane is not being pulled from microvilli during the fast phase and Golgi vesicles only deliver in the slow phase, a second internal reservoir is necessary.

From the above observations, we predict that the slow and fast phases depend on two different reservoirs, one internal and one external, for their primary supplies of cytoskeleton and membrane. We suspect that the slow phase largely relies on the consumption of the apical microvillus membrane, while the fast phase is supported by vesicles and cytoskeletal proteins trafficked directly from a sub-apical compartment (Fig. 7Aii). Therefore in summary, we predict that the slow-to-fast shift may be driven by the sub-apical compartment redirecting vesicle and cytoskeletal transport away from the apical surface as relatively stiff microvilli and into the apical-lateral surface to form a softer, less structured composite. We test these predictions in Figures 7D and 8 next.

As a final set of modeling results, in Figures 7D and 8, we tested the predictive power of an assembly softened scaffold that passes membrane through a sub-apical compartment. First, we evaluated the model against BFA injections in Figure 7D. Then in Figure 8, we evaluated the overall performance of the model against the metrics used in Figures 3, 4, and 5, namely viscoelastic stiffness, wrinkled area accumulation, and furrow length as functions of time. To implement this model, membrane was transported from the Golgi, through the sub-apical compartment, and to the PM as wrinkled membrane with equations 24-29 (Fig. 7A2). The Golgi exported vesicles to the sub-apical compartment for the duration of the slow phase [6], while we chose the sub-apical compartment to exocytose vesicles for the last 45 minutes of cellularization as a free parameter. We initialized the sum of the wrinkled and sub-apical membrane to produce a ∼ 20 µm furrow, because final furrow lengths were measured to be 20 µm after administering BFA injections at ∼ 0 minutes into cellularization [6]. The distribution of membrane area between the two reservoirs was a free parameter. Consistent with Figure 5, the scaffold was softened by the assembly of in-series cytoskeletal elements. Here, cytoskeletal elements were assembled simultaneously with vesicle exocytosis (Eq. 4), but only in the fast-phase – effectively leaving the stiffness of the scaffold unchanged during wrinkled membrane (microvilli) depletion. Additionally, we introduced a minimum value of *R*_*rk*_ termed the elastic floor (*R*_*flr*_) with equation 34. The elastic floor was necessary to produce a similar fast phase velocity under two distinct input conditions: (1) when few vesicles were exocytosed (Fig. 7D, ‘BFA at 0 min.’) and (2) when many vesicles were exocytosed (Fig. 7D, ‘BFA at 33.75 min.’ and ‘no BFA’).

**Figure 8:**
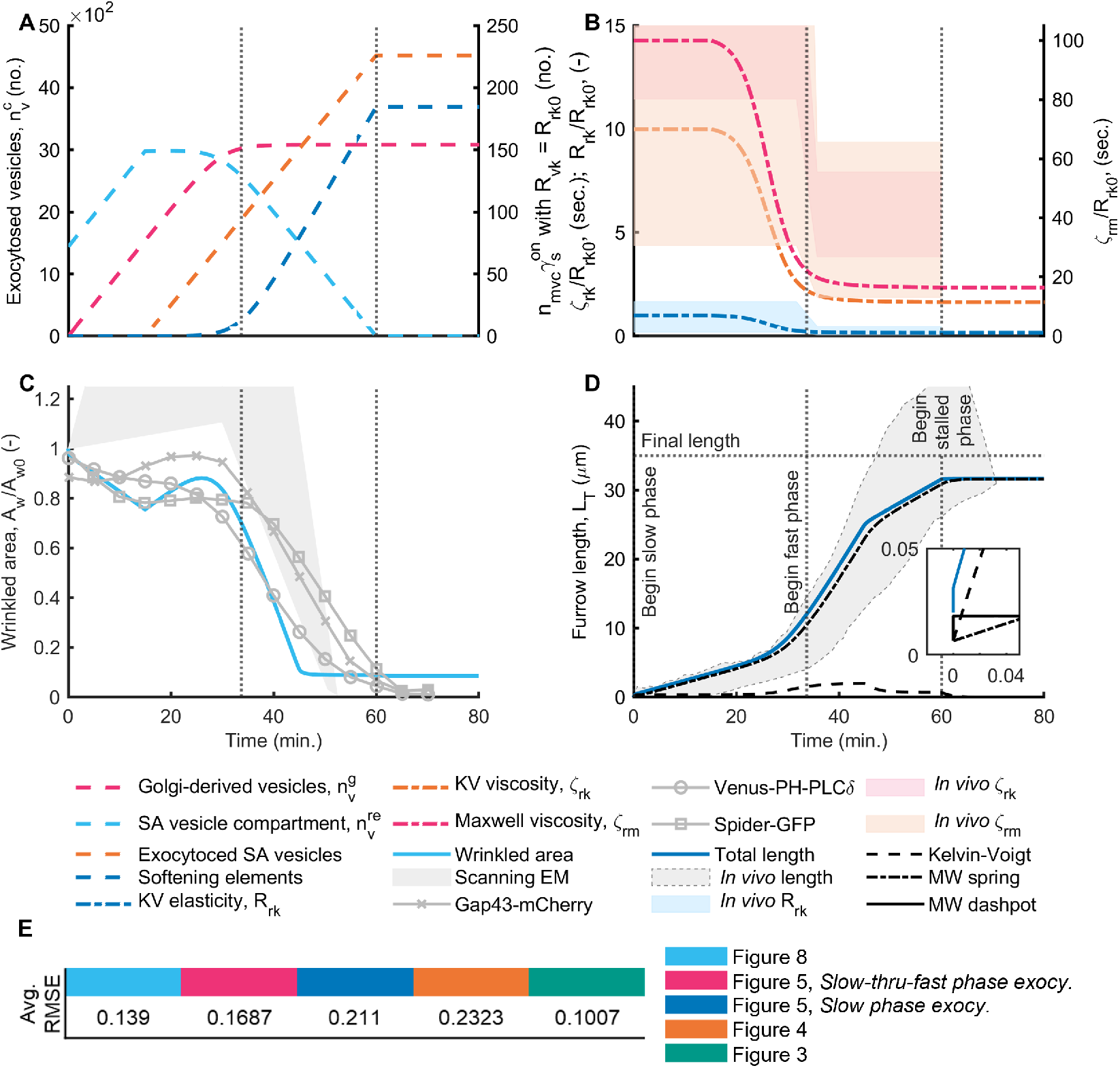
Golgi export through the sub-apical compartment improves the fit of the model to apical arrest and wrinkled membrane data. Figure 8. : (A) The number of vesicles and softening cytoskeletal elements over time. Membrane was transported from the Golgi, through the sub-apical compartment and to the PM with equations 24-29. The duration of Golgi export was minutes, (unitless) for all time, µm, and the duration of sub-apical-to-PM exocytosis was minutes. (B) Change in material properties over time. The membrane-viscoelastic coupling was governed by Equations 30 and 34, where *t* 33.75 minutes, and 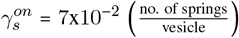. Viscoealstic coefficient data, *In vivo ζ*_*rk*_, *ζ*_*rm*_, and *R*_*rk*_ are from D’Angelo *et al*. [13]. (C) Normalized wrinkled area. Scanning EM data is approximated in Section S1.3 and raw data is from Figard *et al*. [3]. Fluorescence data, Gap43-mCherry, Venus-PH-PLC*δ*, and Spider-GFP are from Figard *et al*. [3]. (D) Time evolution of the furrow length. Length data, *In vivo* length, *Final length, Begin fast phase*, and *Begin stalled phase* are from Figard *et al*. [3]. The gap between the *Final length* and model *Total length* at the end of cellularization is a numerical artifact. All other parameters used in the simulation are listed in Table S1. The apical membrane area, KV spring, and furrow length RMSEs are .181, .155, and .081, respectively (see Figs. S5, S6, and S7 and the caption in Fig. 5). (E) The average total RMSEs are taken from Figs. S5, S6, and S7.

In Figures 4-7C, the transport of membrane and cytoskeletal elements to the PM has been compared to experimental data of composite assembly on the apical surface (Figs. 4 and 5). However, our ODE model is simple in that it does not designate a physical location for wrinkled membrane; wrinkled membrane is just a measure of excess membrane on the PM (Fig. 1E) and its quantity is stored as the non-spatial variable, *A*_*w*_, in equation 22. This spatial ambiguity offers an advantage: the location of vesicle and cytoskeletal delivery along the PM is determined by timing. Specifically, from experimental data [2, 17], transport in the slow phase implies delivery to the apical surface, while transport in the fast phase implies delivery to the apical-lateral surface.

In Figure 7D, we found that the slow and fast velocities as well as the furrow stopping lengths are in agreement with BFA experiments from Figard *et al*. [6]. The slow-to-fast shift occurs even when BFA was injected at 0 minutes into cellularization because initial membrane stored in the sub-apical compartment triggered the velocity shift by transporting vesicles and cytoskeletal elements to the PM 33.75 minutes into cellularization. Further, the furrow stops at the correct length because the initial sum of membrane stored in the wrinkled reservoir and sub-apical compartment was chosen to match the 20 µm found by Figard *et al*. [6]. At the 33.75-minute BFA injection, the furrow stopped at the correct length because by the end of the slow phase the Golgi had already supplied the sub-apical compartment with all of the additional membrane that was needed. After which, the sub-apical compartment transported membrane and cytoskeletal elements to the PM through the end of cellularization. Here, the trafficking of membrane from a sub-apical compartment to the PM is in agreement with Sokac *et al*. [2], Lecuit & Wieschaus [17], and Sisson *et al*. [30].

In Figure 8, we expanded the results of our sub-apical compartment model (Fig. 7D) to evaluate its accuracy in terms of viscoelastic stiffness, wrinkled area, and furrow length (as was used in Figures 3- 5). All parameters and equations used, matched those used in the ‘no BFA’ case of Figure 7D. First, we found that the accumulation of membrane in the sub-apical compartment (Fig. 8A, dashed light blue line) was closely balanced by the influx (dashed magenta line) and efflux (dashed orange line) of vesicles from the Golgi and to the PM, respectively. This agrees with *in vivo* data that found when sub-apical recycling endosome budding (efflux) was inhibited the sub-apical compartment accumulated Golgi-derived membrane [21]. Second, in Figure 8D we found a modest reduction in velocity 45 minutes into cellularization (solid blue line). This occurred because the reservoir of wrinkled area became depleted (Fig. 8C, light blue line), which caused furrow velocity to depend solely on the rate of membrane transport from the sub-apical compartment. Although a slight change of velocity is perceptible at 45 minutes *in vivo* (see Figard *et al*.’s Fig. 2F [3]), in opposition to our model, the *in vivo* change accelerates invagination. Third, we found that the furrow invaginated as a mostly plastic structure. This is illustrated by the Maxwell dashpot (Fig. 8D, dash-dotted black line) having stored most of the length of the furrow (solid blue line), while the Kelvin-Voigt body (dashed black line) and the Maxwell spring (solid black line) contributed less than 5 µm of the total furrow length. This finding agrees with viscoelastic measurements of the cytoskeleton by D’Angelo *et al*. [13] and Doubrovinski *et al*. [14].

As a final measure of accuracy, we found that Figure 8 (an assembly softened scaffold with a sub-apical compartment) is the softening mechanism that most agrees with experimental data [3, 6, 13]. To arrive at this conclusion, we compared the overall average RMSE for each of our models that employed a softening mechanism (RMSE values for Figs. 4, 5, and 8 shown in Fig. 8E). For each mechanism, the overall average RMSE was calculated by averaging the ‘total’ RMSE of viscoelastic stiffness (Fig. S6), wrinkled area (Fig. S5), and furrow length (Fig. S7). Note that Fig. 3 agrees most accurately with experimental data in all categories, but it did not include a softening mechanism and the PM was initialized with enough wrinkled area to complete furrow invagination. Figure 3 strongly suggest scaffold stiffness as the main determinant of invagination velocity. Overall, (1) an assembly-softened scaffold is the most likely driver of the slow-to-fast shift, (2) membrane likely passes through a sub-apical compartment before entering the PM during both slow and fast phases, and (3) more work is needed to understand the exact mapping of membrane trafficking.

## 5. Discussion

Fruit fly embryo cellularization realizes the simultaneous formation of ∼ 6000 new epithelial cells [1, 2]. To accommodate this rapid transformation, the embryo replaces stereotypical cytokinesis with furrow invaginations. The forces driving invagination are unknown due to multiple intersecting mechanisms and molecular components, including motor proteins, microtubules, and F-actin [2].

In this study, we sought to understand the forces governing furrow invagination. Specifically, we were interested in uncovering the regulatory mechanisms that produce the near-instantaneous changes in invagination velocity. Namely, the furrow accelerates midway through cellularization (the slow-to-fast shift) before stalling at the final furrow length (the fast-to-stalled shift) [2, 3, 13, 17]. We used a system of ordinary differential equations to simulate both furrow length and membrane area reservoirs continuously in time. In our model, invagination is driven by motor proteins, while resistive forces come from a viscoelastic cytoskeleton, cytoplasmic drag, and membrane tension [12–15]. We found that the assembly of in series cytoskeletal elements triggers the slow-to-fast shift. Furthermore, we found that an internal sub-apical membrane reservoir likely regulates the trafficking of both cytoskeleton and vesicles to the plasma membrane. This work demonstrates that changes to cell shape may depend on a holistic view of the cell, which includes intimate couplings between the cytoskeleton, membrane reservoirs, trafficking of both, and mechanics.

The slow-to-fast velocity shift can be explained by assembly, and not disassembly, of the scaffold. To accommodate changes in shape, cells store excess membrane in the form of microvilli, where the structure of the microvilli are supported by an underlying cytoskeleton scaffold [3, 6]. When microvilli unwrinkle, the proteins that bind the cytoskeleton to the plasma membrane (PM) can either flow between the phospholipids of the PM or unbind under the increased tension of unwrinkling [22, 27–29]. Disassembly of parallel elements can drive scaffold softening [14]. In our study, we found that scaffold softening due to disassembly, disagreed with experimental data [13]. We found, instead, that the assembly of the in series cytoskeletal elements likely drives the slow-to-fast shift. To reiterate, by in series assembly, we mean the assembly of cytoskeletal elements into the scaffold that support the PM, where each cytoskeletal element (or spring) is added to single-file chain of springs (Fig. 5). This type of assembly has the net effect of softening the overall stiffness of the the collection of springs – or in the case of this paper, the entire scaffold. Our model’s finding is in agreement with *in vivo* data that both vesicles and cytoskeletal elements are transported to the PM for the duration of cellularization [16–19]. Furthermore, we found that the majority of deformation was stored in the Maxwell dashpot, implying that the deformation is plastic (irreversible) [13, 26]. Finally, this result challenges the hypothesis that membrane scarcity is sole governor of invagination velocity in the fast phase [16]. This result may have important implications for other cell types, like neutrophils, which cyclically extend and retract microvillus membrane to engulf foreign debris [23].

A sub-apical membrane reservoir likely regulates the trafficking of the cytoskeleton-membrane composite to the plasma membrane. In place of direct Golgi-to-PM transport of vesicles, cells sometimes favor reservoirs like the recycling endosome to temporarily store and then direct excess membrane into the PM [21]. Further, it has been shown that Golgi-derived vesicles accumulate in a sub-apical reservoir during cellularization, however, the function of sub-apical transport to the plasma membrane is not well understood [21]. We found that a sub-apical compartment was sufficient to replicate *in vivo* invagination behaviors both under normal cellular conditions as well as when the Golgi-derived vesicles were inhibited [6]. However, more work is needed to develop the exact transport pathway between the Golgi, sub-apical compartment, and the PM. Despite the fact that our simulated furrow velocity matched *in vivo* results when Golgi-derived vesicles were inhibited, our wrinkled area failed to replicate *in vivo* data that shows an immediate drop of wrinkled membrane area at the start of the slow phase (shown in Fig. S8D, grey line with diamonds) [6]. Transport from the sub-apical compartment to the PM in the fast phase implies that vesicles and cytoskeletal elements are directed away from the microvillus apical surface (during the slow phase) and towards the relatively flat apical-lateral domain of the furrow. Mechanically, we hypothesize that the slow phase is slow because there is a high density of microvilli, which requires the motor proteins to overcome a higher resulting sliding viscosity between the PM and the proteins that couple the scaffold to the PM (Fig. 5A). Yet, we hypothesize, the fast phase is fast because apical-lateral insertion of composite does not form microvilli and is thus less resistant to expansion. Rationalizing from the perspective of the cell’s efficient use of resources, microvilli are used as a temporary store of excess membrane [2, 3]; during the fast phase, membrane transported directly into the furrow does not require the additional expense of microvilli formation because it is used immediately. The cause of sub-apical redirection to the apical-lateral furrow domain is not well understood, although one possible mechanism may rely on tension-sensitive receptors in the plasma membrane signaling for additional membrane [31–34]. This is supported by Lecuit & Wieschaus’s [17] finding that in the fast phase, microvillus membrane stops being pulled into the furrow, which would create an increased membrane tension along the relatively microvilli-less apical-lateral surface.

Membrane depletion can cause premature furrow stall, but is unlikely to be the mechanism that sets the final furrow stopping length of 35 µm. To complete furrow invagination, the furrow consumes excess membrane stored in a microvillus reservoir [3, 17]. However, at the start of cellularization only about ∼ 50% of the necessary membrane is available in the microvilli; the remainder is provided through vesicle transport [3]. We found that disruption to vesicle transport, resulted in premature stopping lengths of furrows that agreed with *in vivo* findings [6]. Despite our observations, we hypothesize that loss of membrane does not stop the furrow at the end of invagination. This is because the cell continues to consume membrane in order to close its basal end [2]. Furthermore, we extend our hypothesis by proposing that the final furrow length is most likely caused by motor proteins reaching the end of the inverted basket of microtubules, thus preventing motor proteins from continuing to drive the invagination inward. Finally, we propose that the cytokinesis-like constriction described by Krueger *et al*. [5] and Zhang *et al*. [35], becomes the driving force of membrane consumption as the basal end is closed.

In summary, the mechanics of furrow invagination may hold important information as to how the scaffold and membrane interact [2]. However, the vitelline membrane limits access to testing detailed mechanical properties of the furrow [2]. Our model attempts to unite temporally correlated events such as the quantity of apical wrinkled membrane, the underlying mesoscale mechanism of scaffold softening, Brefeldin-A inhibited Golgi, the stoppage of microvillus membrane flow into the furrow in the fast phase, final furrow length, as well as the slow-to-fast and fast-to-stalled transitions. To achieve this task, we proposed a continuum viscoelastic model that governs furrow length dynamics, where necessary membrane additions are supplied by internal sub-apical and external PM reservoirs. However, additional work is needed. Our model does not explore cell shape change from hexagonal to circular at the tip of the invaginating furrow, which too, is temporally aligned with the slow-to-fast shift. Parsimony suggests that the mechanism driving this transformation should be related, if not shared, by the slow-to-fast transition. Our Golgi to sub-apical to compartment to PM pathway is compatible with several prior experimental reports in the literature, including that a that a Rab8/exocyst-dependent pathway of membrane delivery to the subapical furrow surface is necessary in late cellularization [17, 36–38]. However, our model only begins to address how different reservoirs (internal versus external) contribute to different cell surface mechanics.

## 6. Methods

### 6.1 Model geometry

Our model describes how the furrow length increases over time. Thus, we use a system of ordinary differential equations (ODEs) to keep track of the furrow length, given in section 6.6. We approximate the geometry of the invaginating membrane as a cylinder, where the radius of the newly forming cell, *r*_*c*_, is approximated to be constant (Fig. 1E) [2, 4]. In our model, the length of the furrow is equal to the length of the cylinder. The invaginating furrow pulls excess wrinkled membrane from the top of the cylinder down the cylinder walls, to enclose the nucleus and form a new cell (Fig. 1E) [3, 6]. Therefore, our model cylinder has a closed top and open bottom [4, 17]. All parameters used in this model are in Table S1. If other parameters were used in the simulations, it is stated in the corresponding Figure caption.

### 6.2. Force balance

The length of the furrow is governed by a balance of forces acting on the basal tip (Figs. 1B and D). Instead of keeping track of the force felt over the entire circumference of the basal end, *F*, we use the force per circumferential length of the cell, 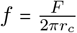, with units 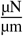. The force balance is given by

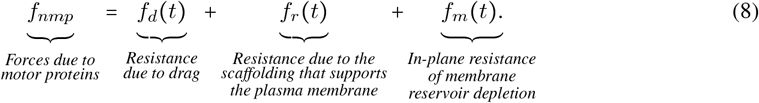

We assume that the net forces exerted by the motor proteins, *f*_*nmp*_, drive the inward invagination of the furrow [2, 16]. We elaborate on the constituents of the net forces in Section 6.2.1. All the other forces resist invagination. The drag force, *f*_*d*_ (see section 6.2.3), accounts for the resistance imposed by a viscous medium on the lengthening furrow. The portion of the scaffold that stabilizes the wrinkled membrane on the apical surface deforms as the furrow lengthens [3, 4, 6], offering an additional resistance, *f*_*r*_, to furrow invagination (see section 6.2.2). Finally, the membrane resists the deformation through in-plane membrane tension, *f*_*m*_, (see section 6.2.4), which is expected to rise rapidly when the external membrane reservoir is depleted and all wrinkled membrane flattens.

#### 6.2.1. Net motor protein forces

The apical surface of the furrow experiences a net tensile force, that is suspected to originate from the plus-end-directed microtubule protein, Pav-KLP [2, 16, 39, 40]. For simplicity, we use a constant force to represent the forces exerted by the motor proteins in all simulations unless noted otherwise explicitly, given by,

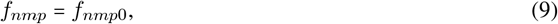

where *f*_*nmp*0_ is the single parameter fit in the slow phase of Figure 3 to ensure the slow phase velocity matches experimental data [6] and is shown in Table S1.

To replicate the increase in tension by oscillatory loading with a magnetic bead [13] (Fig. 6C), we used a square wave,

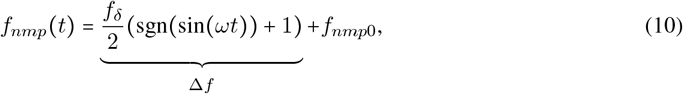

where the oscillation frequency is *ω*, and the maximum change in amplitude is, *f*_*δ*_. To increase the tensile load without oscillations we used,

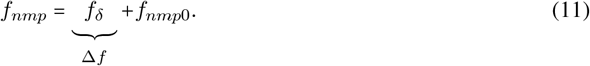

#### 6.2.2. Resistance induced by the cytoskeleton

The tensile resistance induced by the cytoskeleton is modeled as a function of both stored (elastic) and lost (viscous) energy. We represent cytoskeletal tension with a Burger-type viscoelastic body based on experimental findings [13, 14, 41]. Burger-type viscoelastic bodies are composed of a Kelvin-Voigt and Maxwell body connected in series (Figs. 1D and S1A). The Burger body has the advantage of being more general than an isolated fluid or solid model because it deforms with viscous plasticity as well as delayed and instantaneous elasticity. We incorporated the Maxwell spring to represent the instantaneous deformation (or “slack”) found in other microvillus networks [41].

Elements in series necessarily experience the same force, while those in parallel share the load (Fig. S1A). Breaking the Burger body into series elements, each of the sub-elements experience an identical force,

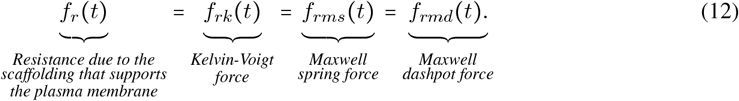

The Kelvin-Voigt element has delayed elasticity because the spring, *R*_*rk*_, is positioned in parallel with the dashpot, *ζ*_*rk*_. The force in this element is given by,

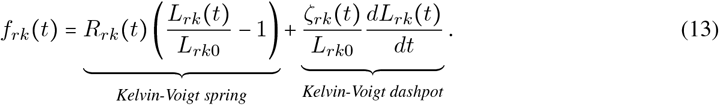

Force applied to the Maxwell spring and dashpot are, respectively, *f*_*rms*_ and *f*_*rmd*_. The spring element, with coefficient *R*_*rm*_, experiences instantaneous elasticity,

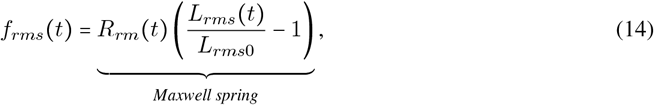

and the dashpot, with coefficient ζ_*rm*_, experiences time-dependent plastic deformation,

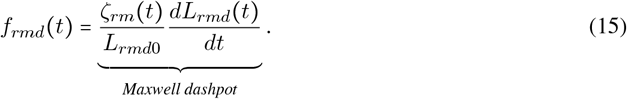

The time-dependent and initial lengths of the spring and dashpot are, respectively, *L*_*rms*_, *L*_*rms*0_, *L*_*rmd*_ and *L*_*rmd*0_. Methods on deriving equations 12, 13, 14, and 15 can be found in Findley *et al*. [24].

##### Elastic-viscous coupling

Viscous coefficients were determined to have a positive, linear scaling with the elastic coefficient of the Kelvin-Voigt body [13]. Hence, we propose

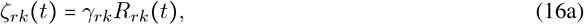

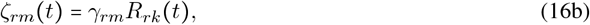

where *γ*_*rk*_ and *γ*_*rm*_ are set to *in vivo* values (Table S1). In section 6.5, we describe how the spring coefficient of the Kelvin-Voigt body is coupled to the exocytosis of vesicles.

##### Slack dynamics

The modulus of elasticity for slack is adapted from the near-instantaneous lengthening of neutrophil lamellipodia during the first 5% of strain [41]. It is defined as,

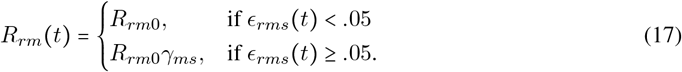

The sudden stiffness encountered upon slack run out is replicated by scaling the Maxwell elastic stiffness by *γ*_*ms*_ (Table S1). The Maxwell spring strain is 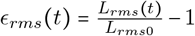1. Neutrophil lamellipodia are suitable representations of the *Drosophila* embryo’s slack strain because they both use microvilli as a membrane reservoir.

#### 6.2.3. Resistance due to drag

The drag force accounts for the friction arising from the invaginating furrow sliding through a viscous medium. We divide the drag force defined in Rangamani *et al*. [25] by 2*πr*_*c*_ to get the force per circumferential length of the furrow,

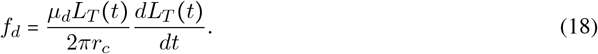

where *µ*_*d*_ is the viscus coefficient and *L*_*T*_ is the total furrow length.

#### 6.2.4. Resistance due to in-plane, membrane tension

In Brefeldin-A experiments, when the microvillus membrane reservoir is restricted, furrow invagination stops at a shortened length and the microvilli flatten on the apical surface prematurely [6]. Importantly, there is also no evidence of membrane rupture. With the preceding evidence, we hypothesize that in-plane membrane tension resists the force of the motor proteins, such that furrow invagination halts when the wrinkled membrane reservoir is exhausted. Therefore, in our model, wrinkled membrane area (*A*_*w*_) depletion, increases in-plane membrane tension as,

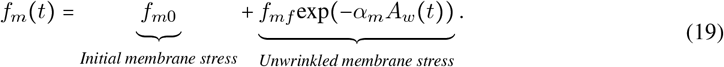

We assume tension from membrane bending is much less than tension from the other stresses in equation 8 and therefore set the initial membrane tension, *f*_*m*0_, equal to zero. The tension from wrinkled membrane area depletion is scaled with *f*_*mf*_ and *α*_*m*_, as *A*_*w*_ → 0. In section 6.3 we give a formal definition for the wrinkled area and how it decays as a function of the furrow length.

### 6.3. Cell membrane area

In section 6.2.4, we proposed how depletion of wrinkled membrane area induces a resistance to furrow invagination. Microvilli unwrinkle as furrow length increases (Fig. 1E) [2, 6, 17]. Furthermore, if the furrow does not lengthen, the microvilli do not deplete [2, 6]. Loss of wrinkled area limits furrow length because it neutralizes the force of the motors (see equations 8 and 19). In this section, we elaborate on how the membrane is distributed in the forming cell and how it behaves during furrow invagination.

In the model, the total cell membrane area, *A*_*c*_, is given by (Fig. 1E),

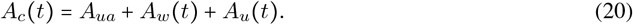

Wrinkled membrane, *A*_*w*_ (Eq. 22), may be enter the furrow when (1) exocytosis of vesicles is directed to the furrow surface or (2) when the KV elastic elements of the Burger body are strained and contribute to furrow length while only consuming a fraction of the wrinkled area. All unwrinkled membrane,

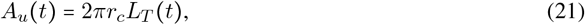

is stored in the furrow (along the axial direction of the cylinder forming the new cell in Fig. 1E). Together, wrinkled and unwrinkled membrane are the only two dynamic states of area in the plasma membrane, *A*_*f*_ (*t*) = *A*_*w*_ (*t*) + *A*_*u*_ (*t*). After the wrinkled membrane area is depleted, the apical surface only keeps an unwrinkled surface with area 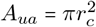

Based on experimental observations [17], we assume that new membrane enters the system in a wrinkled state through vesicle addition, 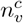. Thus, the wrinkled area is given by,

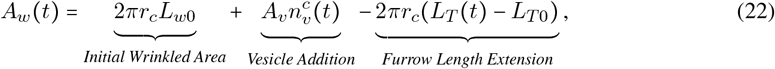

where vesicle radius, vesicle surface area, and initial furrow length are respectively, 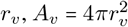 and *L*_*T* 0_. *L*_*w*0_ is the initial length of wrinkled membrane. Therefore, the initial membrane length of the furrow is, *L*_*f*0_ *L*_*w*0_ *L*_*T* 0_. Substituting terms into equation 20 and reducing, results in

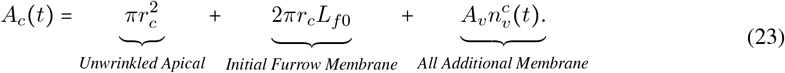

### 6.4. Vesicle sources

Next, we investigate two possible sources of vesicles, the Golgi apparatus (Section 6.4.1) and the sub-apical compartment (Section 6.4.2). In Figures 3-7C, we assume that all vesicles come from the Golgi apparatus and then we introduce the sub-apical compartment in Figures 7D and 8.

#### 6.4.1. The Golgi as a source

In our model, we assume that the Golgi acts as a source of vesicles, and that enough vesicles are released to provide the furrow with all of the additional membrane needed to reach an invagination length of 35 µm. The rate of vesicle transport from the Golgi is given by,

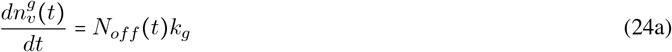

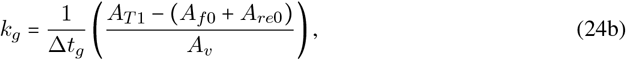

where the initial condition, 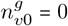. The sum of membrane supplied by the Golgi is then, *A*_*T* 1_−(*A*_*f*0_+*A*_*re*0_), where *A*_*re*0_ = 2*πr*_*c*_*L*_*re*0_ is the initial total area of the vesicles in the sub-apical compartment, and the input parameter, *L*_*re*0_, is the length-equivalent of membrane stored in the sub-apical reservoir at *t* = 0. From *in vivo* data [6], the sub-apical reservoir and wrinkled membrane store 20 µm of membrane at the start of the slow phase, therefore, *L*_re0_ = 20 − *L*_*re*0 *f*0_. When there is no initial membrane in the sub-apical reservoir, or when the Golgi exocytoses directly to the furrow, *L*_*re*0_ *=* 0. Here, *A*_*T* 1_, is the total area needed to complete furrow invagination (Table S1), and, *A*_*f*0_, is the initial, dynamic membrane in the furrow, stored in both wrinkled and unwrinkled states (as described in section 6.3). Golgi export stops after a duration of Δ*t* = 30_*g*_ minutes of cellularization, based on *in vivo* data (Fig. 7A) [6]. *N*_*off*_ = 1 when *t* < *t*_*bfa*_ and 0 otherwise.

When simulating the the Brefeldin-A test (see Section 4.6.2), we set

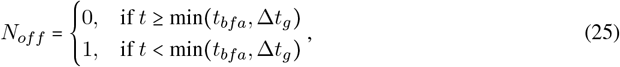

where the injection time is the input parameter, *t*_*bfa*_. BFA only stops Golgi-derived vesicles, which are assumed to begin exporting at *t* = 0, thus making Δ*t*_*g*_ an effective export end time.

#### 6.4.2 Sub-apical compartment

When the sub-apical (SA) compartment is included in the membrane pathway (Fig.7A2), the amount of stored area, *A*_*re*_ (*t*), increases as a function of Golgi vesicles transported to the SA compartment and decreases as a function of vesicle exocytosis from the SA compartment to the PM surface, 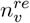,

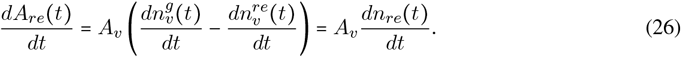

For convenience, rather than tracking SA compartment area, we solve for the area-equivalent number of vesicles stored in the SA compartment, *n*_*re*_.

The rate of vesicle transport from the SA to the plasma membrane is given by,

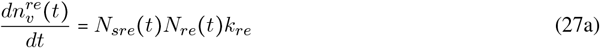

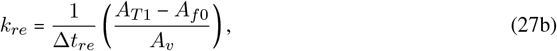

where, *A*_*f*0_, is the initial membrane in the furrow, and Δ*t*_*re*_ is the duration of vesicle delivery. Exocytosis stops when the SA reservoir is depleted, as,

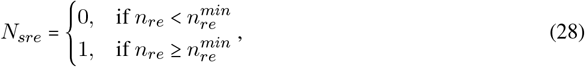

and begins when,

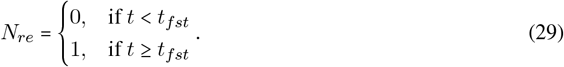

Here, the minimum fraction of vesicles required to continue exocytosis, 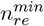, is defined in Table S1, and the SA exocytosis start time, *t*_*fst*_, was chosen to fit fluorescent area data curves in Figure 8.

### 6.5. The effects of vesicles and invagination on the resistive forces of the scaffold

Next, we investigate how vesicle addition, and motor protein tension change the properties of the underlying scaffold and, therefore, the dynamics of furrow invagination. Instead of modeling individual cytoskeletal filaments and their crosslinking proteins, we lump the mesoscale mechanisms with a continuum representation of the scaffold material properties (Fig. 1D, Section 6.2.2) and assume that the constitutive properties are represented by the parameters of the Burger body, namely, the spring constants, *R*, and dashpot viscosities, *ζ* (Fig. S1A). The resulting, lumped constitutive properties are governed by the properties of its individual units (the cytoskeletal filaments), its connectivity with itself (cytoskeletal filament density and crosslinking proteins), and the geometry of those connections (cytoskeleton filament orientations) (Figs. 6A and 7A; S1) [14, 27–29]. Cytoplasmic viscosity is excluded here because it does not model scaffold deformation and assembly, rather it represents the bulk movement of the plasma membrane-bound scaffold through a medium.

#### 6.5.1. Temporal relation between vesicle exocytosis and scaffold assembly

Here, we introduce equations that allow us to control the timing of viscoelastic changes to the scaffold as a function of vesicle exocytosis. For this, we introduce a new variable *n*_*mvc*_ which coordinates the timing of scaffold assembly as a function of vesicle exocytosis. In the first type of coupling, used in Figures 5 (*Slow-thru-fast phase exocytosis*); 7C; 6D; and 8, vesicles begin affecting viscoelasticity at *t*_*on*_ minutes and are never disabled,

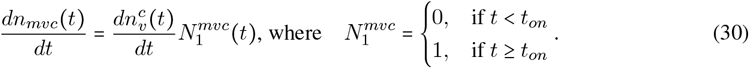

The superscript, *c*, indicates these equations can be used for either Golgi or the SA trafficking.

In the alternate coupling (Figs. 5 (*Slow phase exocytosis*) and 7B), we explore a delay between vesicle exocytosis and viscoelastic changes to the PM. Because there is a supported coupling between the quantity of microvillus membrane and the quantity of F-actin [6], the rate of scaffold assembly is still proportional to the rate of exocytosis,

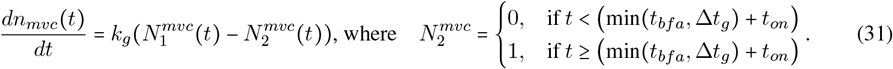

Here, 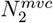 is subtracted from 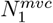 to set the duration of the vesicle exocytosis (Tables S1, S2).

#### 6.5.2 Fit against experimental data

In Figure 3, we used experimental data to dictate when and by how much the Kelvin-Voigt elasticity changes. It does not assume a mechanism of change. It was leveraged to (1) fit a net motor protein force (Eq. 9) to the slow phase velocity, (2) demonstrate that a continuum, viscoelastic model replicates furrow behavior, and (3) determine the magnitude of change to the KV elasticity, *γ*_*R*_ (Table S1), needed to produce the fast phase velocity.

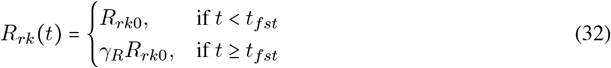

#### 6.5.3 Assembly-softened scaffold

In this module, we assume that during the fast phase cytoskeletal elements are assembled into the scaffold in series (Figs. 5; 6D; 7B,C; S1A-C). Further, we couple the transport of vesicles and cytoskeletal elements to the plasma membrane based on experimental findings [16, 18, 19]. The derivation of how series elements affect overall elasticity is in the *Supplementary Material* section S1.1. The rate of change of the Kelvin-Voigt spring is,

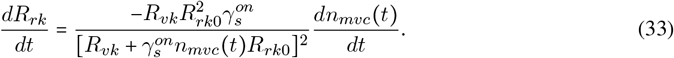

The fraction of series elements contributed by each vesicle is fitted as, 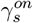. Without an experimentally measured stiffness of individual cytoskeletal springs, *R*_*vk*_, we set each disassembled spring equal to the initial stiffness of the total scaffold, *R*_*rk*0_, measured by D’Angelo *et al*. [13] (Table S1).

When we use vesicles from the SA compartment to soften the plasma membrane (Figs. 7D, 8), the BFA test only works if there is a softening floor, below which additional vesicles have no effect on the membrane. Physically, we suspect this is because after a wrinkle has flattened, the membrane becomes a load path parallel to the cytoskeleton, such that the elastic elements beneath it no longer contribute to the cortical strain. Thus, the cortical-membrane reaches a minimum softness, *R*_*flr*_ (Table S1),

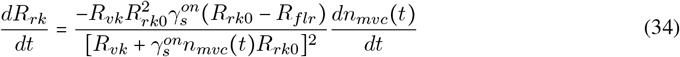

#### 6.5.4. Disassembly-softened scaffold

In Figures 4 and 6E, F, we assume that parallel assembly stiffens the scaffold, while tension-driven tension disassembles and softens the scaffold (Fig. S1A-C). Cytoskeletal and vesicle transport are coupled by,

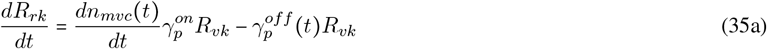

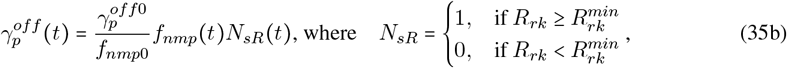

where 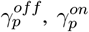, and 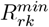 are the disassembly rate of parallel cytoskeletal elements due to an applied tension on the scaffold, the assembly rate of cytoskeletal elements as a function of vesicle and cytoskeleton transport to the plasma membrane, and the minimum elastic stiffness of the Kelvin-Voigt body, respectively. Note that the rate of change of cortical elasticity decreases with increased applied tension, *f*_*nmp*_, and passes through the fitted force-unbinding rate point, 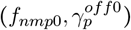.

### 6.6. Governing equations and numerical implementation

To examine how the furrow length increases over time, we derive the following governing equations

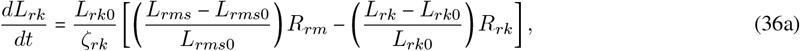

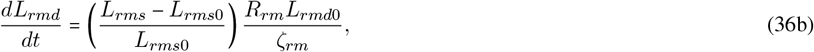

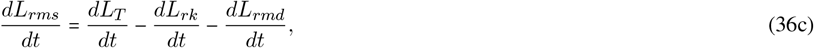

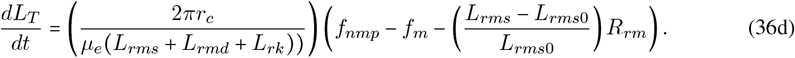

Total furrow velocity, 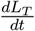 (Eq. 36d), is derived in the supplement section S1.2 from the force balance equation 8. Equation 36a comes from the center equality in equation 15, *f*_*rms*_ = *f*_*rk*_, followed by rearranging terms. Equation 36b comes from the right-most equality in equation 15, *f* = *f*_*rms rmd*_ followed by rearranging terms. Finally, equation 36c is the time-derivative of length-continuity, *L*_*T*_ = *L*_*rk*_ + *L*_*rms*_ + *L*_*rmd*_. This system of equations are numerically solved in Matlab2023a with ODE solver, ode15s. The value for the initial total length is provided in Table S1, with 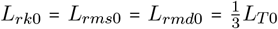 Equations governing the net motor force, in-plane membrane force, viscoelastic coefficients are found in sections 6.2.1, 6.2.4, 6.2.2 (*Slack dynamics* and *Elastic-viscous coupling*), respectively. All other parameter values are listed in Table S1.

#### 6.6.1. Smoothing “on/off” functions

Introducing sudden transitions in the ODE system with conditional logic (if-statements, in the case), can introduce numerical errors. To minimize errors, we use the continuous and smooth hyperbolic tangent for all piecewise functions,

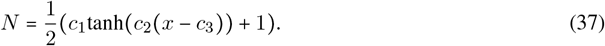

The transition from zero to one, or one to zero is set *c*_1_ = 1, −1, respectively. The slope of the transition is *c*_2_, the independent variable is *x*, and the position along the x-axis that initiates the transition is *c*_3_ (Table S2).

#### 6.6.2. Oscillatory net motor force

For simplicity, we used the *square* function in Matlab to implement the square wave oscillation of the net motor force defined in eq. 10, 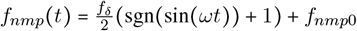.

## S1 Supplementary Material

### S1.1 Cytoskeletal assembly

#### S1.1.1 Derivation of assembly softening

Viscoelastic elements are summed in-series as,

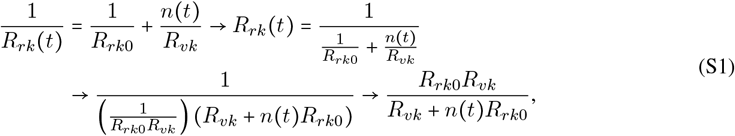

where *n* is the number of assembled springs. For a mechanical representation of the springs, see Figures 5A and S1 [42]. Taking the first derivative in time produces Equation 33.

#### S1.1.2 Proof that series springs can only either soften or have no effect

To prove that springs summed in-series can only reduce (soften) or produce no change in the elastic coefficient, the below limits were evaluated, where *n* is assumed to be greater than zero.

**Limit 1:** We first evaluate the effects of adding a single, infinitely stiff spring (*R*_*vk*_ → ∞) as a series element,

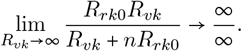

Applying L’Hopital’s rule, we get,

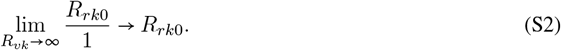

Therefore, *R*_*rk*_ → *R n*_*rk*0_, and this results in no change to the elastic coefficient of the system. Because drops out of the equation after applying L’Hopital’s rule, we can conclude that *R*_*rk*_ →*R*_*rk*0_ is true for any number of series springs.

**Limit 2:** Next, we evaluate the effects of adding an infinite number of infinitesimally soft springs to the system (*n* → ∞, *R*_*vk*_ → 0) as series elements,

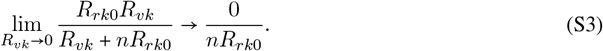

Then, taking the limit as *n* goes to infinity, we get,

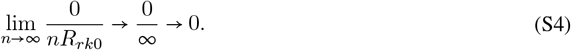

Therefore, *R*_*rk*_ → 0, and this results in an infinitesimally small (soft) elastic coefficient of the system.

Limits S.S3 and S.S4 demonstrate that this is true for both a finite and infinite number of series springs.

**Limit 3:** Finally, we evaluate the effects of adding an infinite number of springs with a stiffness coefficient equal to the initial system stiffness (*n* → ∞, *R*_*vk*_ → *R*_*rk*0_) as series elements,

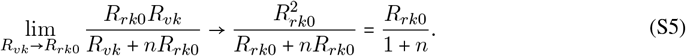

Then, taking the limit as *n* goes to infinity, we get,

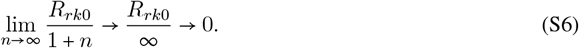

Therefore, as *n*→∞ and *R*_*vk*_ → *R*_*rk*0_, *R*_*rk*_ → 0, the elastic coefficient of the system becomes infinitesimally small (soft) as 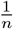 (Fig. S1).

#### S1.1.3 Derivation of assembly softening with a lower bound

To introduce a non-zero, softening, lower bound (*R*_*flr*_) on series-spring addition, we constrain Equation S1,

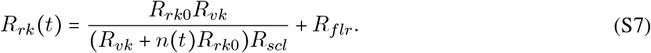

The y-intercept is kept unchanged by setting, *R*_*rk*_(*t* = 0) = *R*_*rk*0_, which results in 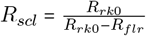 and,

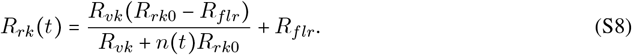

Finally, taking the first derivative in time produces Equation 34.

#### S1.1.4 Derivation of assembly stiffening

Viscoelastic elements are summed in parallel in the form of equation S9 [42]. For a mechanical representation of the springs, see Figures 4A and S1.

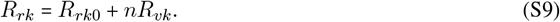

#### S1.1.5 Proof that parallel springs can only stiffen or have no effect

To prove that springs summed in-parallel can only increase (stiffen) or produce no change in the elastic coefficient, the below limits were evaluated, where *n* is assumed to be greater than zero.

**Limit 1:** We first evaluate the effects of adding a single, infinitely stiff spring (*R*_*vk*_ → ∞) as a parallel element,

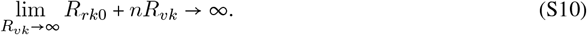

Therefore, *R*_*rk*_ → ∞, resulting in an infinitely large (stiff) elastic coefficient of the system. This is true for both a finite and infinite number of series springs.

(*R*_*vk*_ → 0) as a parallel element,

**Limit 2:** Next, we evaluate the effects of adding a single, infinitesimally soft spring to the system

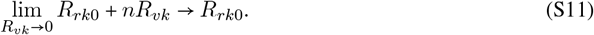

Therefore, *R*_*rk*_ → *R*_*rk*0_, resulting in no change to the elastic coefficient of the system.

**Limit 3:** Finally, we evaluate the effects of adding an infinite number of springs with a stiffness coefficient equal to the initial system stiffness (*n* → ∞, *R*_*vk*_ → *R*_*rk*0_) as parallel elements,

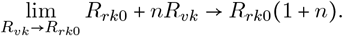

Then, taking the limit as *n* goes to infinity, we get,

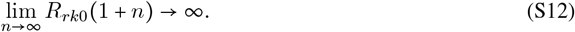

Therefore, as *n* → ∞ and *R*_*vk*_ → *R*_*rk*0_, → *R*_*rk*_, the elastic coefficient of the system becomes infinitely large (stiff) as *n* (Fig. S1).

### S1.2 Total furrow velocity

Combining equations 8, 9, 14, 18 and 19 we get,

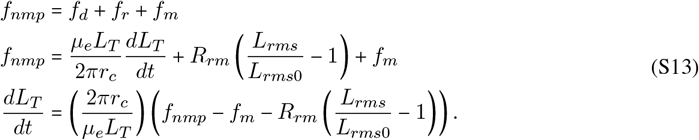

Our choice to represent actin scaffold resistance, *f*_*r*_, with the Maxwell spring, *R*_*rm*_, reduces algebraic errors. This choice takes advantage of the property that a series of viscoelastic elements share a single load path, and therefore experience the same stress. Equation 12 states this directly.

### S1.3 Estimated microvillus reservoir area from scanning EM data

To estimate the accumulation of apical membrane area from scanning EM data, we used *in vivo* data of furrow velocity, initial membrane stored in microvilli, and the final amount of membrane needed to form a 35 µm furrow. The slow and fast phase velocities are, respectively, 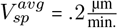 with a range of 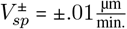, and 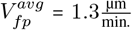. with a range of 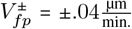. [3]. The necessary final amount of membrane area is, *A*_*T*1_ (Table S1) [3]. The estimates of membrane at the start of cellularization reflect all “finger-like” and “hand-like” microvillus structures. Respectively, theses are 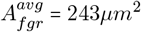 with a range of 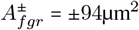 and 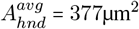with a range of 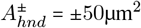[3].

The area-contributions of vesicles needed at the start of cellularization are,

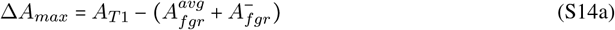

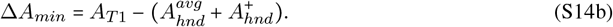

The corresponding velocities of apical area accumulation in the slow phase are, respectively,

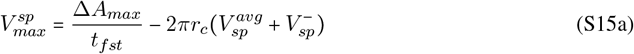

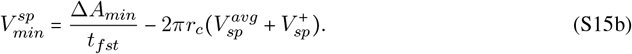

To show the widest spread of apical membrane decay in the fast phase, the Δ*A*_*max*_ case decays the slowest and the Δ*A*_*min*_ case decays the fastest,

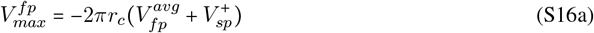

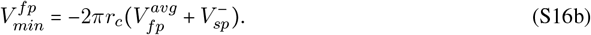

Plotting each of these velocities in their respective, slow and fast phases produces an upper and lower bound of apical membrane area. Plotted values are normalized by the apical membrane at *t* = 0 minutes.

### S1.4 Supplementary figures and tables

**Table S1:** Constant parameters. *Superscripts (a),(c),(h) were chosen to prevent numerical instability, but have little impact on the results. (b) The Maxwell elasticity was assumed to match the measured Kelvin-Voigt spring. (d) R*_*flr*_ equals *R*_*rk*_ in the fast phase of Fig. 3. (e),(f) were chosen to be as minimize response time without causing numerical errors. (g) Measured in the center of a *Drosophila* embryo. (i) Chosen to achieve the correct fast phase velocity in Fig. 3.

| Symbol | Description | Value | Units | References |
| --- | --- | --- | --- | --- |
| $A_{T1}$ | Final membrane area | 580 | $\mu m^2$ | [3] |
| $r_c$ | Cell radius | 2.7 | $\mu m$ | [3, 14] |
| $r_v$ | Vesicle radius | .075 | $\mu m$ | [43] |
| $L_{T0}$ | Furrow length at t = 0 min | .02 | $\mu m$ | [17]<sup>a</sup> |
| $L_{T1}$ | Furrow length at t = 60 min | 35 | $\mu m$ | [3, 14, 17] |
| $f_{m0}$ | Initial membrane force | 0 | $\frac{\mu N}{\mu m}$ | Section 6.2.4 |
| $f_{mf}$ | Membrane tension scaling factor | 100 | $\frac{\mu N}{\mu m}$ | Section 6.2.4 |
| $f_{nmp0}$ | Net motor force | $5.15 \times 10^{-3}$ | $\frac{\mu N}{\mu m}$ | Fitted |
| $f_\delta$ | Stress test: additional load | $10.35 \times 10^{-4}$ | $\frac{\mu N}{\mu m}$ | [13], Section 6.2.1 |
| $\omega$ | Stress test: osc. angular freq. | 120 | $\frac{1}{s}$ | [13] |
| $R_{rk0}$ | Kelvin-Voigt spring coeff., init. est. | $9.8 \times 10^{-5}$ | $\frac{\mu N}{\mu m}$ | [13] |
| $R_{rm0}$ | Maxwell spring coeff., init. est. | $3 \times 10^{-5}$ | $\frac{\mu N}{\mu m}$ | [13], Fitted<sup>b</sup> |
| $R_{rk}^{min}$ | Minimum softness (unbinding) | $5 \times 10^{-6}$ | $\frac{\mu N}{\mu m}$ | Fitted<sup>c</sup> |
| $R_{flr}$ | Minimum softness (sliding) | $1.568 \times 10^{-5}$ | $\frac{\mu N}{\mu m}$ | Fitted<sup>d</sup> |
| $\gamma_{rk}$ | Elastic-viscous coupler (Kelvin-Voigt) | 10 | $s$ | [13] |
| $\gamma_{rm}$ | Elastic-viscous coupler (Maxwell) | 100 | $s$ | [13] |
| $\gamma_{ms}$ | ”Slack” scalar | 100 | # | Fitted<sup>e</sup> |
| $\gamma_R$ | Slow-to-fast scalar | .16 | # | Fitted<sup>i</sup> |
| $\alpha_m$ | Membrane tension sensitivity | .5 | $\frac{1}{\mu m^2}$ | Fitted<sup>f</sup> |
| $\mu_d$ | Cytosol viscous coeff. | $15 \times 10^{-6}$ | $\frac{\mu N * s}{\mu m^2}$ | [14]<sup>g</sup> |
| $n_{re}^{min}$ | Minimum RE compartment size | $10^{-3}$ | # | Fitted<sup>h</sup> |
| $t_{slw}$ | Start of slow-phase | 0 | $min$ | [3, 17] |
| $t_{fst}$ | Start of fast-phase | 30 | $min$ | [3, 17] |
| $t_{std}$ | Start of stalled-phase | 60 | $min$ | [3, 17] |

**Table S2:**
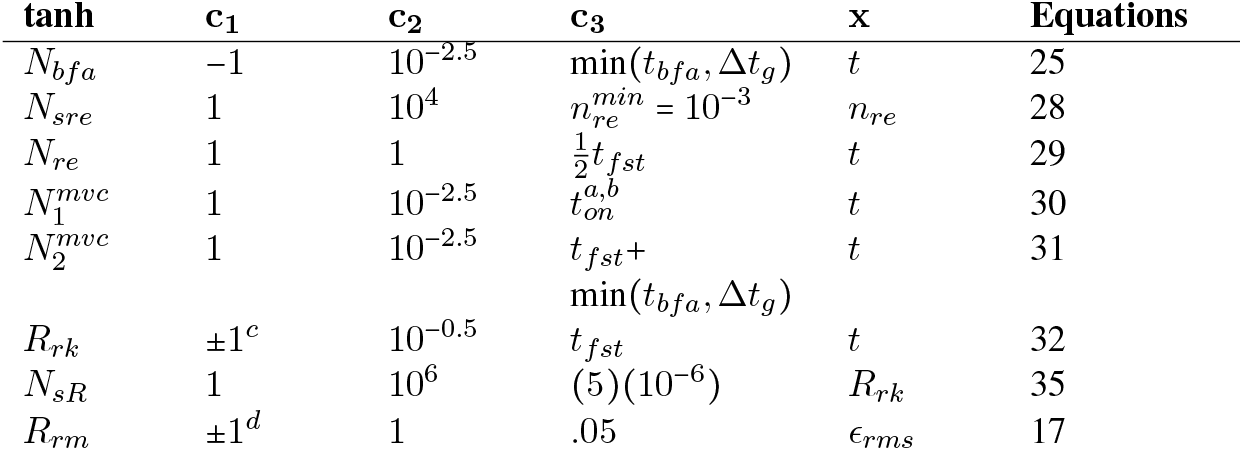
Hyperbolic tangent parameters for smoothing function Eq. 37. (a) minutes for the disassembly-softened scaffold. (b) minutes for the assembly-softened scaffold. (c) Two hyperbolic tangent curves are used simultaneously. The first, *c*_1_ = −1, decays the slow phase elasticity to *R*_*rk*_ = .16*R*_*rk*0_, while the second, *c*_1_ = +1, scales the fast phase elasticity to *R*_*rm*_ = 0 (d) Two hyperbolic tangent curves are used simultaneously. The first,, decays the slow phase elasticity to, while the second, *c*_1_ = +1, scales the fast phase elasticity to *R*_*rm*_ = 100*R*_*rm*0_.

**Figure S1:**
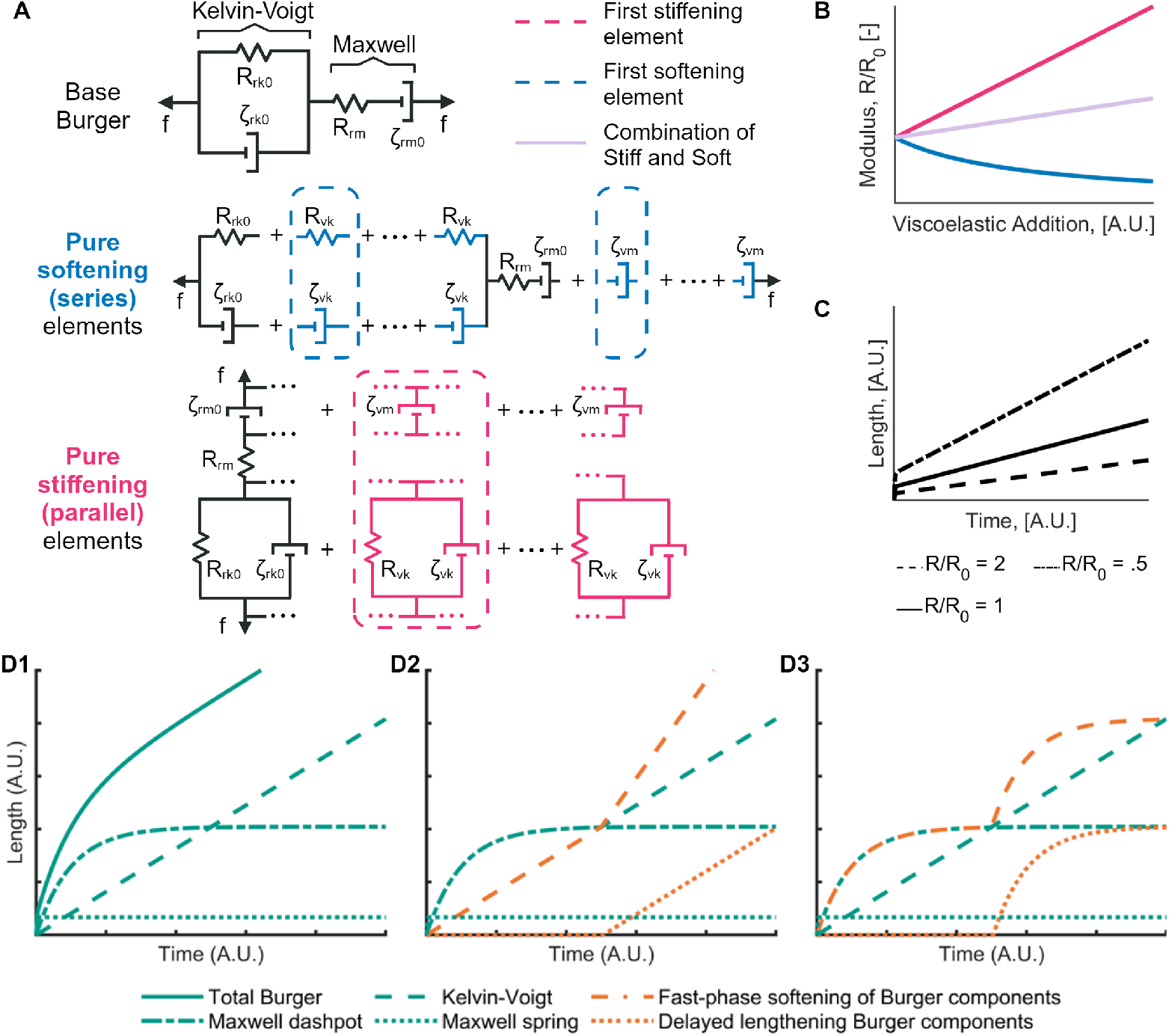
A geometric representation of the scaffold with series and parallel spring additions. (A) The *Base Burger* body lumps the network cytoskeletal filaments beneath the plasma membrane into a single continuous body. Additional cytoskeletal elements affect the Kelvin-Voigt and Maxwell dashpots with Equations 16A and B. (B) The summation of series cytoskeletal elements (*Pure Softening* decays the lumped elasticity as, 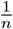, where *n* is the number of assembled springs). The summation of parallel cytoskeletal elements (*Pure Stiffening*) increases the lumped elasticity as, *n*. (C) Furrow velocity negatively correlates with lumped elasticity, *R*, where *R*_0_ is the initial elastic coefficient. (D1) Length of the Burger body under constant applied stress and constant viscoelastic coefficients. Lengths of Burger components (the Maxwell dashpot, Maxwell spring, or Kelvin-Voigt body) with non-constant viscoelastic coefficients in the Maxwell dashpot (D2) and the Kelvin-Voigt body (D3).

**Figure S2:**
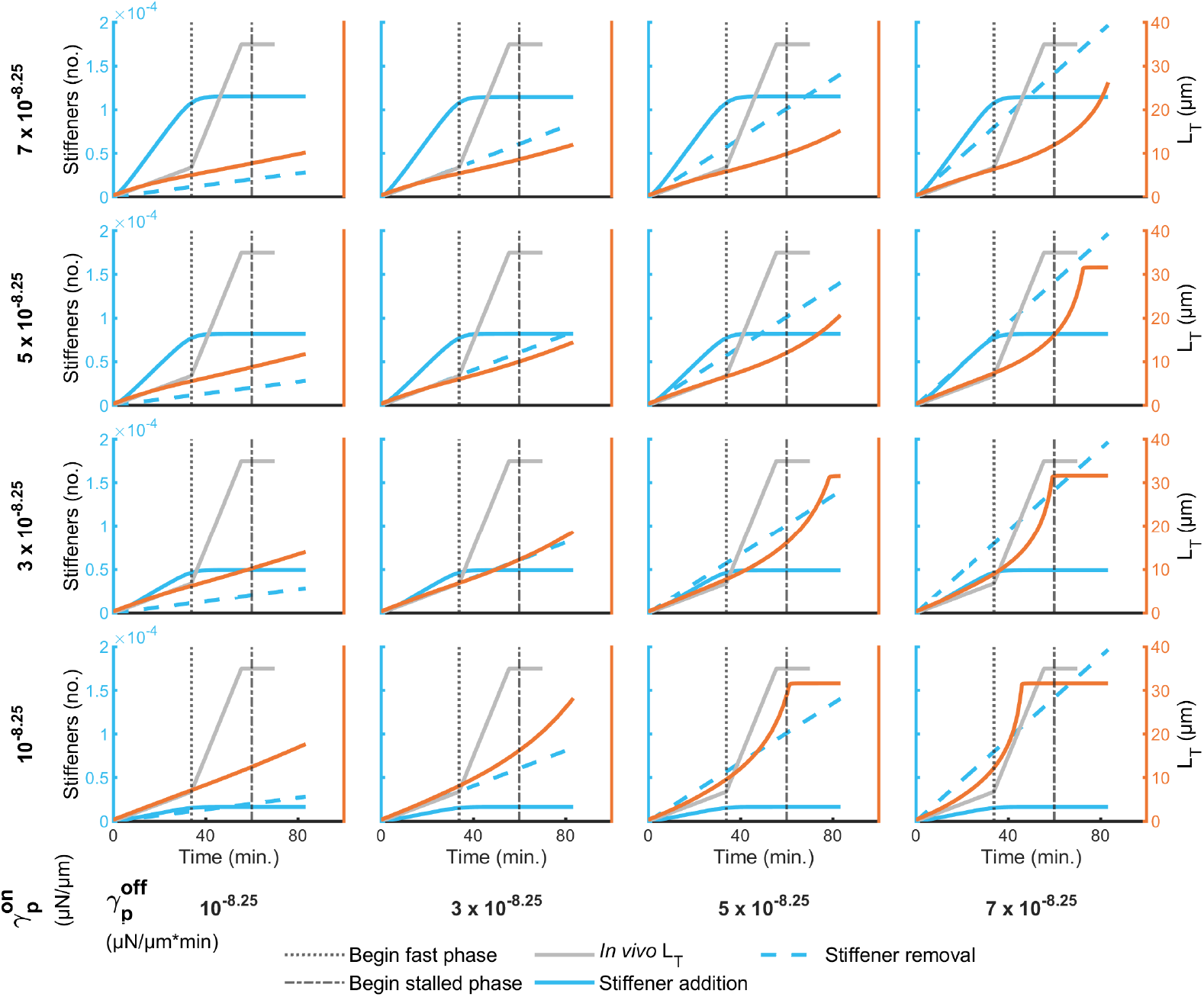
Furrow behavior as a function of varied assembly and disassembly rates of stiffening, viscoelastic elements. Furrow length (orange line) was affected by the competition between the addition of stiffening elements (solid blue line, Fig. S1B, *Pure Stiffening*) and tension-dependent removal of stiffening elements (dashed blue line) in the scaffold disassembly model (Figs. 4B, S1). Viscoelastic properties were stiffened as a function of Golgi exocytosis to the PM domain, where membrane was added only in the slow phase using Eq. 24, with *N*_*off*_ = 1 for all time and *L*_*re*0_ = 0. The timing and velocities of the slow, fast, and stalled phases of the furrow were determined by sweeping the assembly (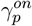, vertical direction) and disassembly 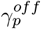, horizontal direction) rates between 10^−8.25^ and 7 × 10^−8.25^. Viscoelastic properties were governed by Eqs. 30 and 35, where *f*_*nmp*_ = *f*_*nmp*0_, and *t*_*on*_ = 0 minutes (Table S2) The gap between the *Final length* and model *Total length* (orange line) at the end of cellularization is a numerical artifact. *In vivo L*_*T*_, *Begin fast phase*, and *Begin stalled phase* are from Figard *et al*. [3]. All other parameters are listed in Table S1.

**Figure S3:**
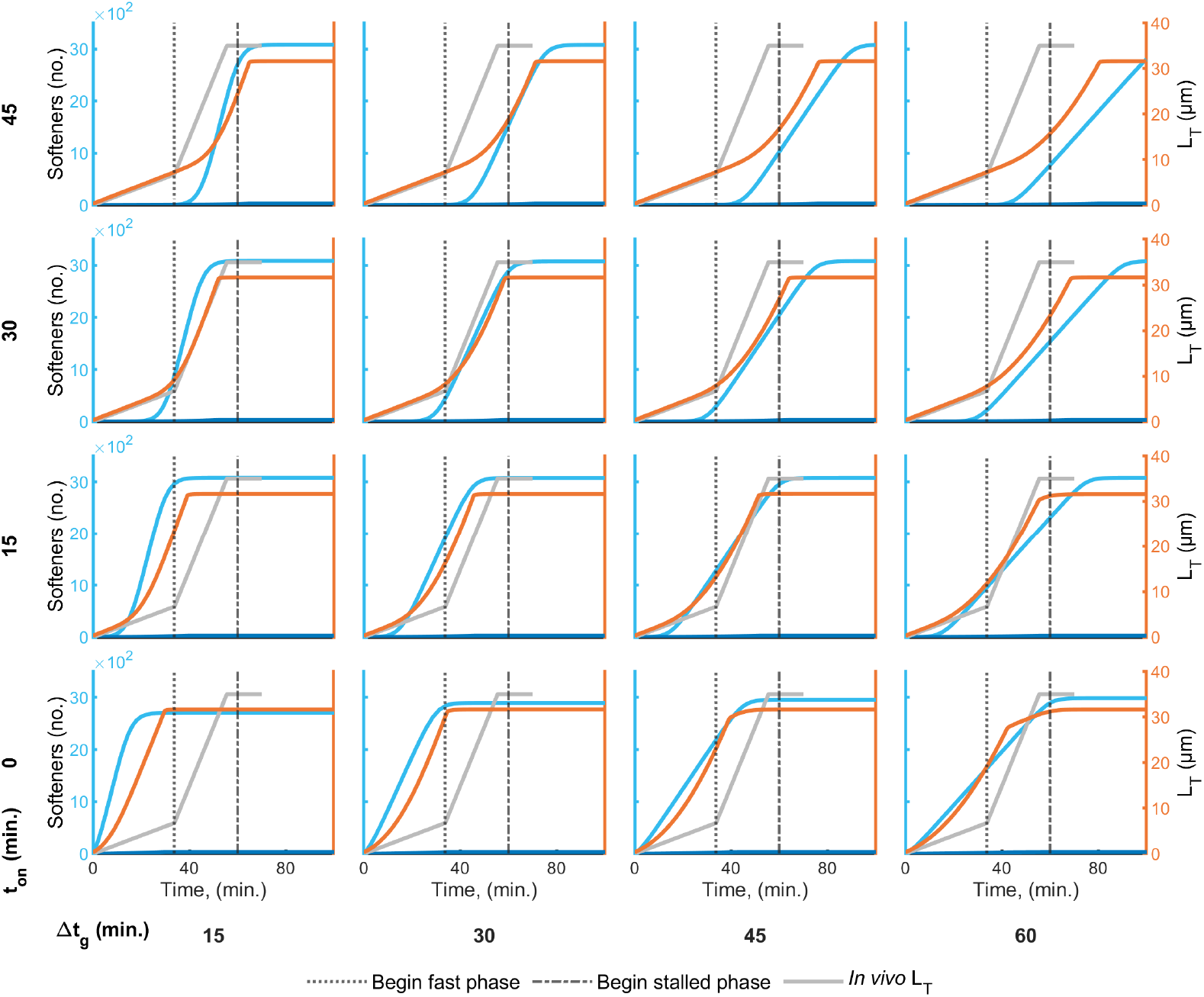
Furrow behavior when the timing and duration of assembly-softened cytoskeleton addition are varied. Furrow length (orange) was affected by the timing of softening elements (blue, Fig. S1A, *Pure Softening*) added to the assmebly-softened scaffold (Figs. 5, S1). Viscoelastic properties were softened as a function Golgi exocytosis to the PM domain, where membrane was added only in the slow phase using Eq.24, with *N*_*off*_ = 1 for all time and *L*_*re*0_ = 0. The timing between exocytosis and viscoelastic softening was determined by sweeping a delayed, softening start time between 0 and 45 min. (*t*_*on*_, vertical direction), and the duration of Golgi exocytosis (Δ*t*_*g*_, horizontal direction). Viscoelastic properties are governed by Eqs. 31 and 33, where 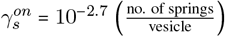. The gap between the *Final length* and model *Total length* (blue lines) at the end of cellularization is a numerical artifact. *In vivo L*_*T*_, *Begin fast phase*, and *Begin stalled phase* are from Figard *et al*. [3]. All other parameters are listed in Table S1.

**Figure S4:**
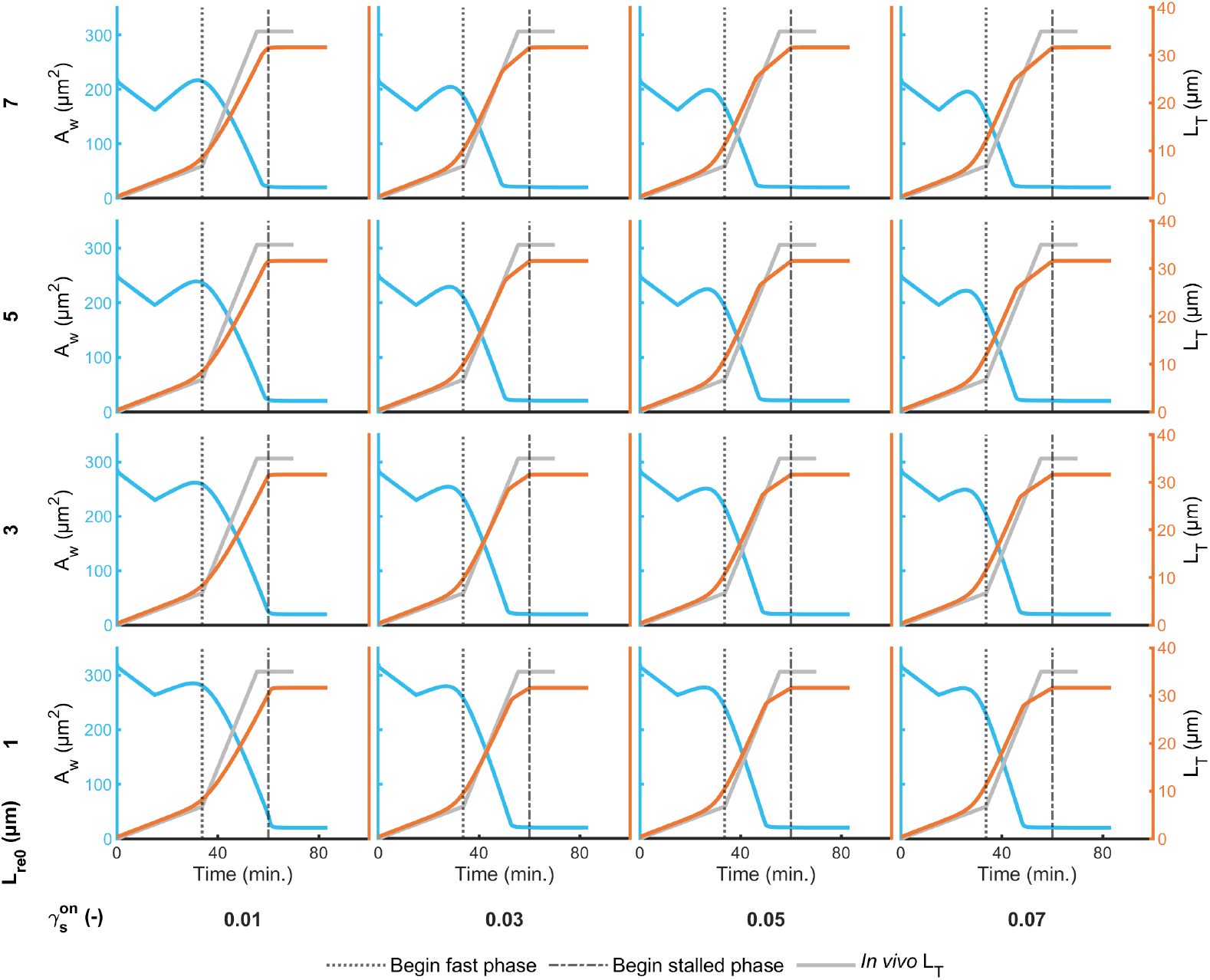
Furrow and wrinkled membrane reservoir behavior as a function of initial sub-apical membrane storage and vesicle-induced softening. Membrane was transported from the Golgi, through the sub-apical reservoir, and into the PM domain with Equations 24 - 28. The duration of Golgi export Δ*t*_*g*_ = 30 was minutes, *N*_*reoff*_ = 1 for all time, and the duration of sub-apical transport to the PM was Δ*t* = 45_*g*_ minutes. The vesicle-cytoskeleton coupling was governed by equations 30 and 34, where *t*_*on*_ = 33.75 min. The timing and velocities of the slow, fast, and stalled phases of the furrow were determined by sweeping the initial length of membrane stored in the sub-apical compartment (*L*_*re*0_, vertical direction) and the softening effectiveness of each vesicle on the scaffold (*γ*^*on*^, horizontal direction), between 1 and 7µm and .01 and .07 (-), respectively. *L*_*re*0_ = 6 µm, and 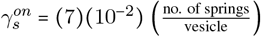. Viscoealstic coefficient data, *In vivo ζ*_*rk*_, *ζ*_*rm*_, and *R*_*rk*_ are from D’Angelo *et al*. [13]. *In vivo* length, *Begin fast phase*, and *Begin stalled phase* are from Figard *et al*. [3]. The gap between the *in vivo L*_*T*_ and the model *L*_*T*_ at the end of cellularization is a numerical artifact. All other parameters are listed in Table S1.

**Figure S5:**
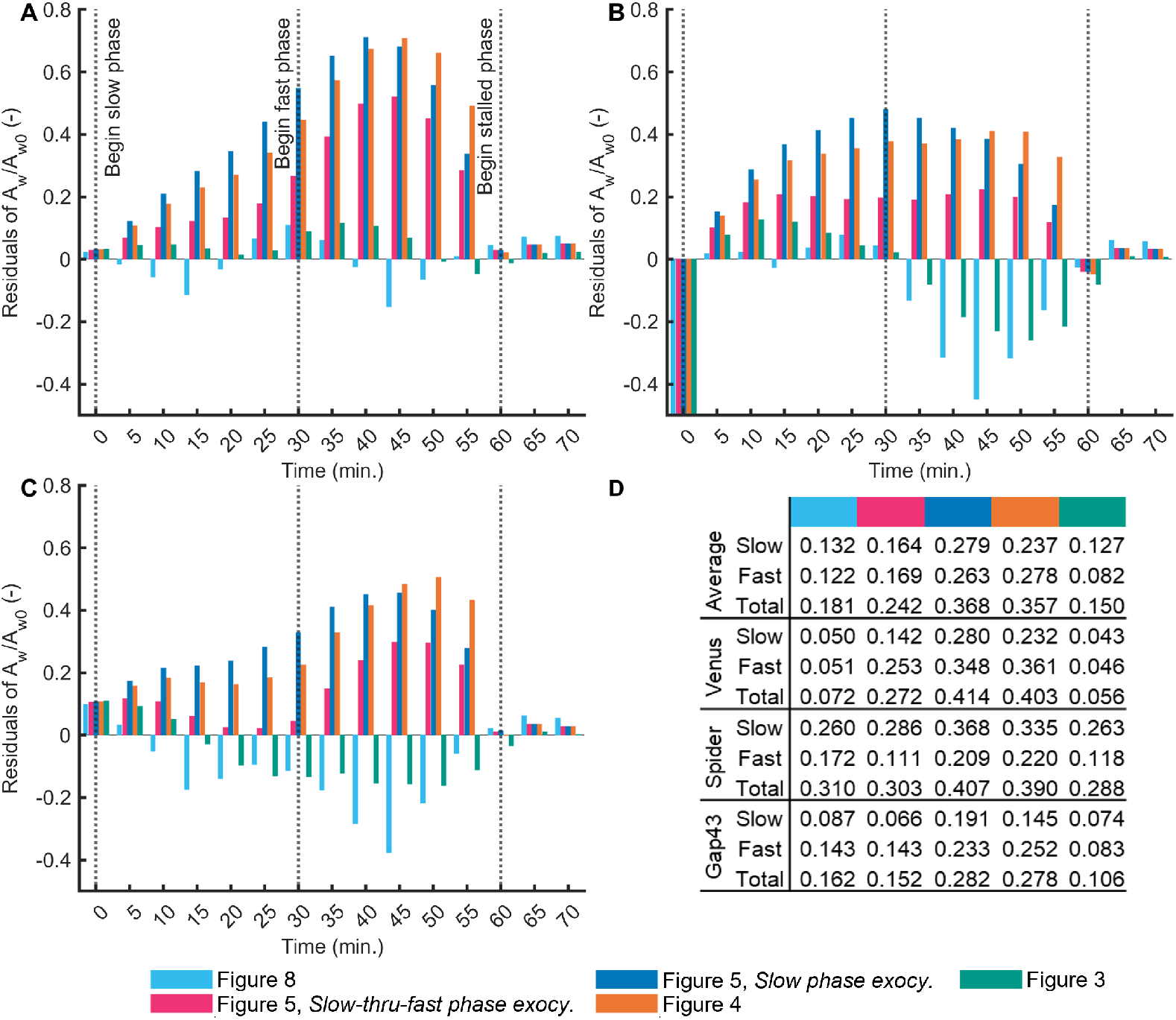
Area residuals. Residual errors between model mechanisms (color coded in the legend) and fluorescent markers Venus (A), Spider (B) and Gap43 (C). (B) Residuals in the 0-min. bin extend to approximately -1. (D) Norms of the residuals from panels (A-C), where ‘Slow’, ‘Fast’ and ‘Total’ evaluated the norms over 0-30 min, 35-65 min, and 0-70 min, respectively. The *Average* cases average the norms of the three fluorescent measurements. I.e. the ‘Average Slow’ value for the Figure 8 case is the average of each ‘Slow’ Figure 8 norm, .132 = (.050 + .260 + .087)/3.

**Figure S6:**
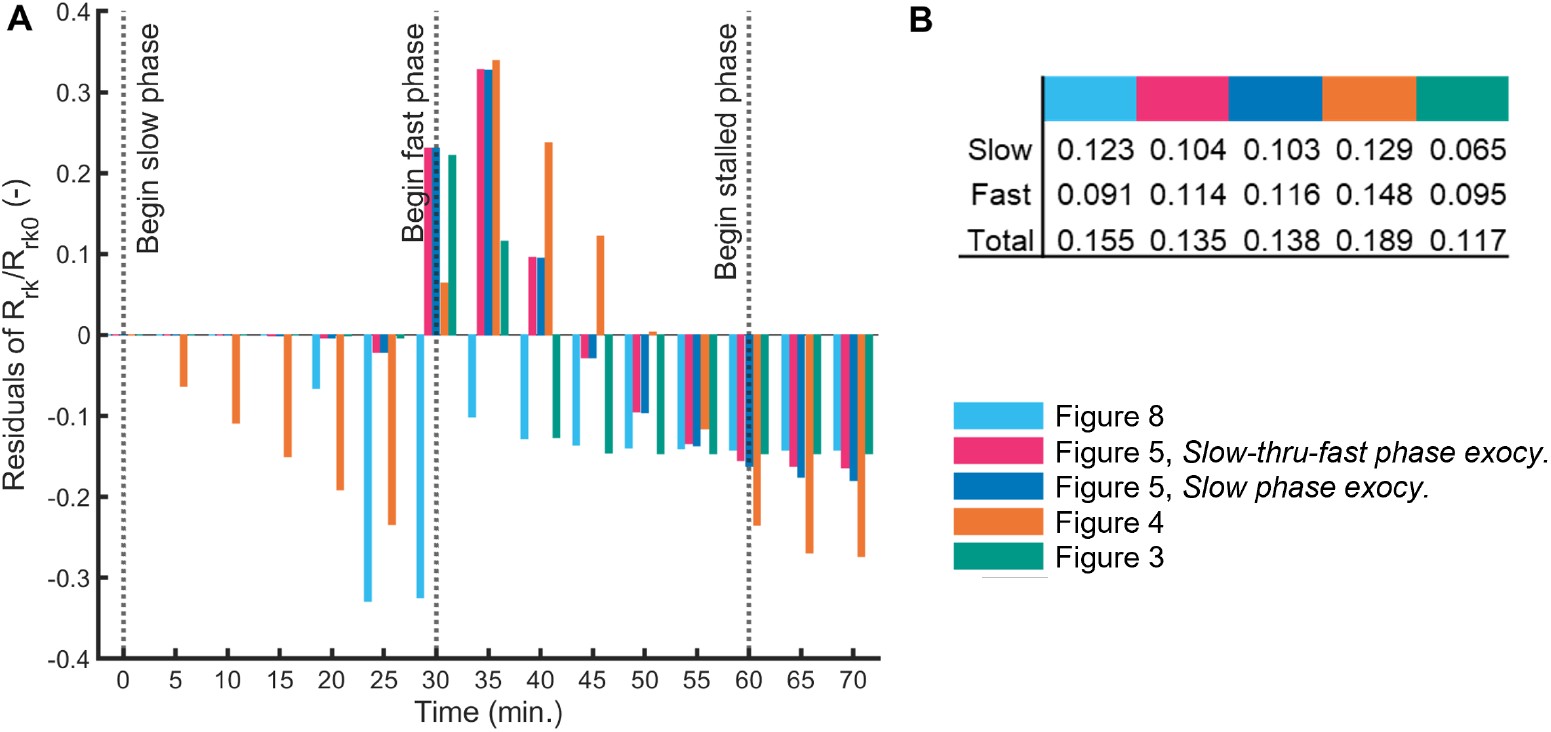
Viscoelastic residuals. (A) Residual errors between model mechanisms (color coded in the legend) and the Kelvin-Voigt elastic coefficient, *R*_*rk*_. (B) Norms of the residuals from panel (A), where ‘Slow’, ‘Fast’ and ‘Total’ evaluated the norms over 0-30 min, 35-65 min, and 0-70 min, respectively.

**Figure S7:**
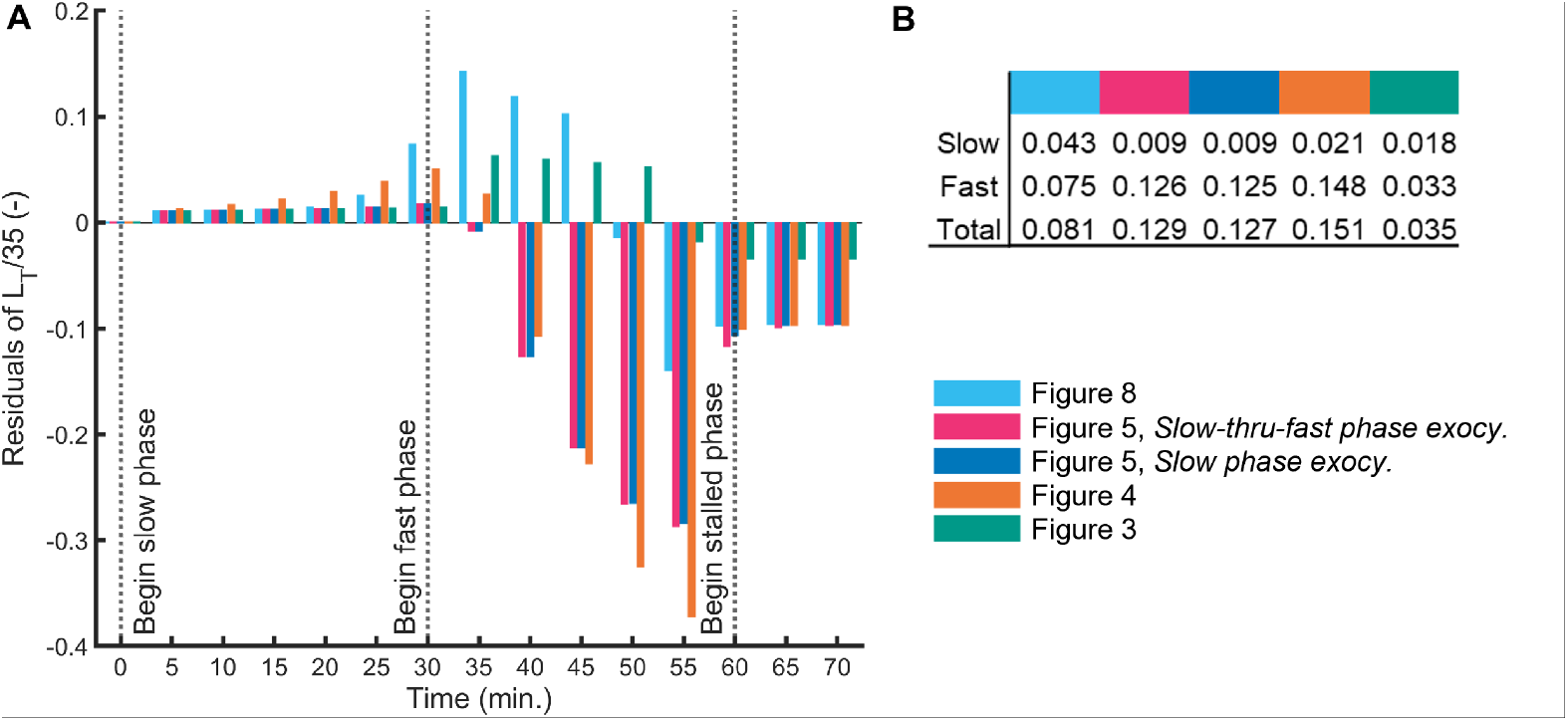
Length residuals. (A) Residual errors between model mechanisms (color coded in the legend) and the *in vivo* furrow length, *L*_*T*_, normalized by the furrow stopping length, 35 µm. (B) Norms of the residuals from panel (A), where ‘Slow’, ‘Fast’ and ‘Total’ evaluated the norms over 0-30 min, 35-65 min, and 0-70 min, respectively.

**Figure S8:**
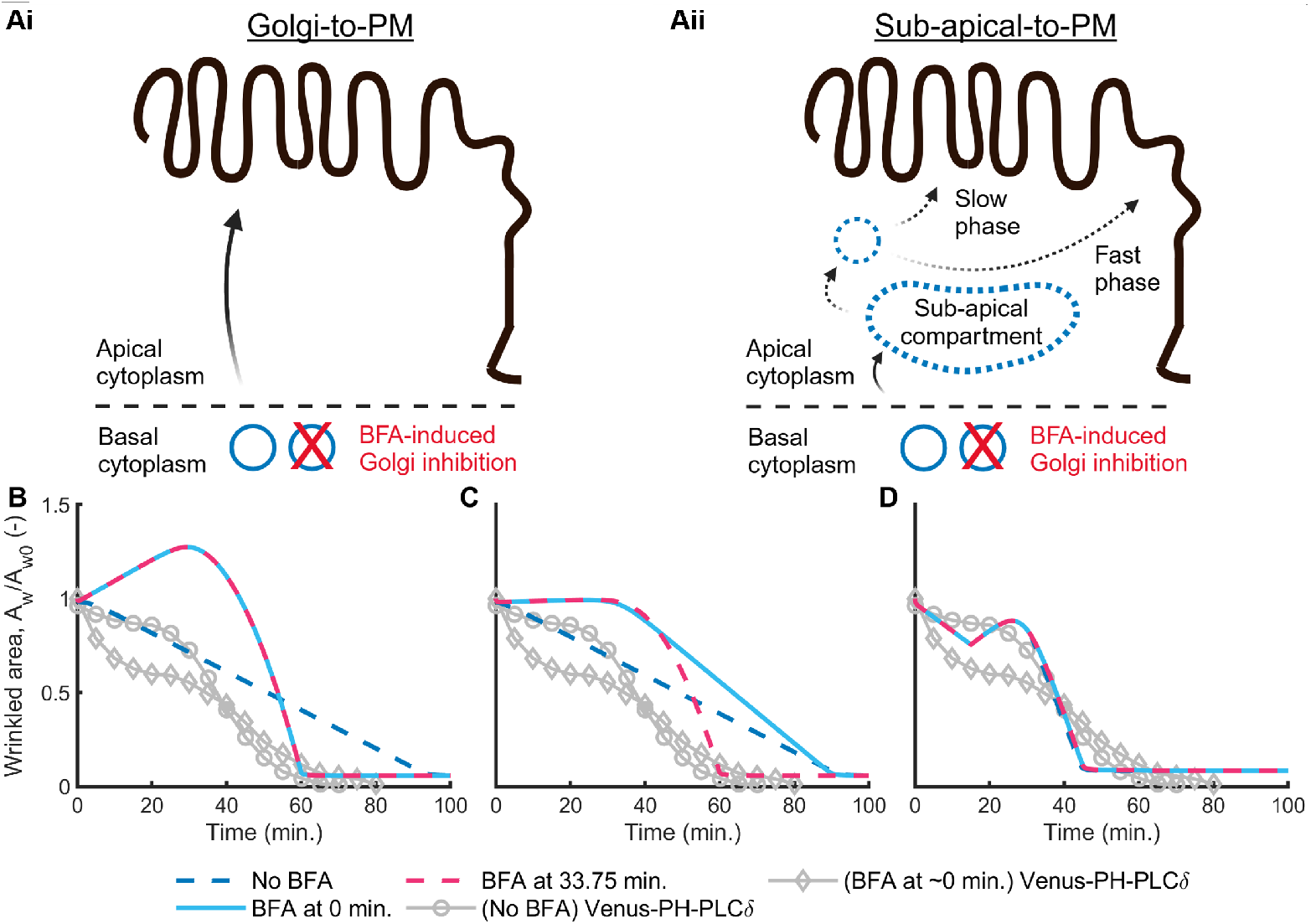
Golgi transport through a sub-apical compartment replicates Brefeldin-A (BFA) results. (Ai, Aii) Golgi-derived vesicles originate in the basal cytoplasm of the embryo and enter the space of the apical cytoplasm. BFA inhibits the formation of Golgi-derived vesicles [6]. (Ai) Golgi-derived vesicles are exocytosed directly to the PM domain. Panels B and C used this pathway. (Aii) Golgi-derived vesicles are first transported to a sub-apical compartment and then sub-apical vesicles are exocytosed to the PM domain. Panel D used this pathway. (B-D) The time evolution of wrinkled area, when Golgi-derived vesicle transport was stopped at *t*_*bfa*_ = 0 and 33.75 minutes into cellularization (Eq. 25). *In vivo* data is from Figard *et al*. [6]. (B, C) Vesicles were trafficked directly from the Golgi, and exocytosed to the PM (panel Ai). (B) This simulation extended the results from Figure 5, *slow phase exocytosis*, in which all vesicles were exocytosed during the slow phase (first 33.75 min.) and viscoelastic changes to the scaffold were delayed by 30 minutes. Excluding *t*_*bfa*_, all other parameters and equations are unchanged. (C) This simulation extended the results from Figure 5, *slow-thru-fast phase exocytosis*, in which vesicles were exocytosed throughout cellularization (60 minutes) and viscoelastic changes to the scaffold take effect only in the fast phase (last 26.25 min.). Excluding *t*_*bfa*_, all other parameters and equations are unchanged. (D) Membrane was transported from the Golgi, through the sub-apical compartment, and into the PM with equations 24-29 (panel Aii). The duration of Golgi export was Δ*t*_*g*_ = 33.75 minutes, *L*_*re*0_ = 6 µm, and the duration of sub-apical-to-PM exocytosis was Δ*t*_*re*_ = 45 minutes. The membrane-viscoelastic coupling was governed by equations 30 and 34, where *t*_*on*_ = 33.75 min., and 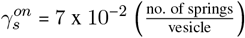. All other parameters are listed in Table S1.

## Notes

### Competing Interest Statement

The authors have declared no competing interest.

## References

1. Foe, V. E. & Alberts, B. M. Studies of nuclear and cytoplasmic behaviour during the five mitotic cycles that precede gastrulation in Drosophila embryogenesis. en. J. Cell Sci. 61, 31–70 (May 1983).

2. Sokac, A. M., Biel, N. & De Renzis, S. Membrane-actin interactions in morphogenesis: Lessons learned from Drosophila cellularization. en. Semin. Cell Dev. Biol. 133, 107–122 (Jan. 2023).

3. Figard, L., Xu, H., Garcia, H. G., Golding, I. & Sokac, A. M. The plasma membrane flattens out to fuel cell-surface growth during Drosophila cellularization. en. Dev. Cell 27, 648–655 (Dec. 2013).

4. Afshar, K., Stuart, B. & Wasserman, S. A. Functional analysis of the Drosophila diaphanous FH protein in early embryonic development. en. Development 127, 1887–1897 (May 2000).

5. Krueger, D., Quinkler, T., Mortensen, S. A., Sachse, C. & De Renzis, S. Cross-linker-mediated regulation of actin network organization controls tissue morphogenesis. en. J. Cell Biol. 218, 2743–2761 (Aug. 2019).

6. Figard, L., Wang, M., Zheng, L., Golding, I. & Sokac, A. M. Membrane Supply and Demand Regulates F-Actin in a Cell Surface Reservoir. en. Dev. Cell 37, 267–278 (May 2016).

7. Lemerle, E. et al. Caveolae and Bin1 form ring-shaped platforms for T-tubule initiation. en. Elife 12 (Apr. 2023).

8. Quinn, C. J. & Dibb, K. M. The origin of T-tubules. en. Elife 12 (June 2023).

9. Klecker, T. & Westermann, B. Pathways shaping the mitochondrial inner membrane. en. Open Biol. 11, 210238 (Dec. 2021).

10. Mahapatra, A., Uysalel, C. & Rangamani, P. The Mechanics and Thermodynamics of Tubule Formation in Biological Membranes. en. J. Membr. Biol. 254, 273–291 (June 2021).

11. Venkatraman, K. et al. Cristae formation is a mechanical buckling event controlled by the inner mito-chondrial membrane lipidome. en. EMBO J. 42, e114054 (Dec. 2023).

12. Mogilner, A. & Manhart, A. Intracellular Fluid Mechanics: Coupling Cytoplasmic Flow with Active Cytoskeletal Gel. Annu. Rev. Fluid Mech. 50, 347–370 (Jan. 2018).

13. D’Angelo, A., Dierkes, K., Carolis, C., Salbreux, G. & Solon, J. In Vivo Force Application Reveals a Fast Tissue Softening and External Friction Increase during Early Embryogenesis. en. Curr. Biol. 29, 1564–1571.e6 (May 2019).

14. Doubrovinski, K., Swan, M., Polyakov, O. & Wieschaus, E. F. Measurement of cortical elasticity in Drosophila melanogaster embryos using ferrofluids. en. Proc. Natl. Acad. Sci. U. S. A. 114, 1051–1056 (Jan. 2017).

15. Shi, Z., Graber, Z. T., Baumgart, T., Stone, H. A. & Cohen, A. E. Cell Membranes Resist Flow. en. Cell 175, 1769–1779.e13 (Dec. 2018).

16. Sommi, P. et al. A mitotic kinesin-6, Pav-KLP, mediates interdependent cortical reorganization and spindle dynamics in Drosophila embryos. en. J. Cell Sci. 123, 1862–1872 (June 2010).

17. Lecuit, T. & Wieschaus, E. Polarized insertion of new membrane from a cytoplasmic reservoir during cleavage of the Drosophila embryo. en. J. Cell Biol. 150, 849–860 (Aug. 2000).

18. Cao, J., Albertson, R., Riggs, B., Field, C. M. & Sullivan, W. Nuf, a Rab11 effector, maintains cytoki-netic furrow integrity by promoting local actin polymerization. en. J. Cell Biol. 182, 301–313 (July 2008).

19. Riggs, B. et al. Actin cytoskeleton remodeling during early Drosophila furrow formation requires recycling endosomal components Nuclear-fallout and Rab11. en. J. Cell Biol. 163, 143–154 (Oct. 2003).

20. Albertson, R., Cao, J., Hsieh, T.-S. & Sullivan, W. Vesicles and actin are targeted to the cleavage furrow via furrow microtubules and the central spindle. en. J. Cell Biol. 181, 777–790 (June 2008).

21. Pelissier, A., Chauvin, J.-P. & Lecuit, T. Trafficking through Rab11 endosomes is required for cellu-larization during Drosophila embryogenesis. en. Curr. Biol. 13, 1848–1857 (Oct. 2003).

22. Borghi, N. & Brochard-Wyart, F. Tether extrusion from red blood cells: integral proteins unbinding from cytoskeleton. en. Biophys. J. 93, 1369–1379 (Aug. 2007).

23. Herant, M., Heinrich, V. & Dembo, M. Mechanics of neutrophil phagocytosis: behavior of the cortical tension. en. J. Cell Sci. 118, 1789–1797 (May 2005).

24. in. Creep and Relaxation of Nonlinear Viscoelastic Materials - With an Introduction to Linear Vis-coelasticity (eds Findley, W. N., Davis, F. A. & Onaran, K.) 50–107 (New York: Dover Publications, 1990).

25. Rangamani, P., Zhang, D., Oster, G. & Shen, A. Q. Lipid tubule growth by osmotic pressure. J. R. Soc. Interface 10, 20130637 (Nov. 2013).

26. Goldner, A. N. et al. Evidence that tissue recoil in the early Drosophila embryo is a passive not active process. en. Mol. Biol. Cell 34, br16 (Sept. 2023).

27. Alvarado, J., Sheinman, M., Sharma, A., MacKintosh, F. C. & Koenderink, G. H. Molecular motors robustly drive active gels to a critically connected state. en. Nat. Phys. 9, 591–597 (Aug. 2013).

28. Lieleg, O., Claessens, M. M. A. & Bausch, A. R. Structure and dynamics of cross-linked actin net-works. en. Soft Matter 6, 218–225 (Jan. 2010).

29. Corominas-Murtra, B. & Petridou, N. I. Viscoelastic Networks: Forming Cells and Tissues. Frontiers in Physics 9 (2021).

30. Sisson, J. C., Field, C., Ventura, R., Royou, A. & Sullivan, W. Lava lamp, a novel peripheral golgi protein, is required for Drosophila melanogaster cellularization. en. J. Cell Biol. 151, 905–918 (Nov. 2000).

31. Figard, L. & Sokac, A. M. A membrane reservoir at the cell surface: unfolding the plasma membrane to fuel cell shape change: Unfolding the plasma membrane to fuel cell shape change. en. Bioarchitecture 4, 39–46 (Mar. 2014).

32. Masters, T. A., Pontes, B., Viasnoff, V., Li, Y. & Gauthier, N. C. Plasma membrane tension orchestrates membrane trafficking, cytoskeletal remodeling, and biochemical signaling during phagocytosis. en. Proc. Natl. Acad. Sci. U. S. A. 110, 11875–11880 (July 2013).

33. Gauthier, N. C., Masters, T. A. & Sheetz, M. P. Mechanical feedback between membrane tension and dynamics. en. Trends Cell Biol. 22, 527–535 (Oct. 2012).

34. Gauthier, N. C., Fardin, M. A., Roca-Cusachs, P. & Sheetz, M. P. Temporary increase in plasma membrane tension coordinates the activation of exocytosis and contraction during cell spreading. en. Proc. Natl. Acad. Sci. U. S. A. 108, 14467–14472 (Aug. 2011).

35. Zhang, Y., Yu, J. C., Jiang, T., Fernandez-Gonzalez, R. & Harris, T. J. C. Collision of Expanding Actin Caps with Actomyosin Borders for Cortical Bending and Mitotic Rounding in a Syncytium. Dev. Cell 45, 551–564.e4 (June 2018).

36. Murthy, M., Teodoro, R. O., Miller, T. P. & Schwarz, T. L. Sec5, a member of the exocyst complex, mediates Drosophila embryo cellularization. en. Development 137, 2773–2783 (Aug. 2010).

37. Holly, R. M., Mavor, L. M., Zuo, Z. & Blankenship, J. T. A rapid, membrane-dependent pathway directs furrow formation through RalA in the early Drosophila embryo. en. Development 142, 2316– 2328 (July 2015).

38. Mavor, L. M. et al. Rab8 directs furrow ingression and membrane addition during epithelial formation in Drosophila melanogaster. en. Development 143, 892–903 (Mar. 2016).

39. He, B., Martin, A. & Wieschaus, E. Flow-dependent myosin recruitment during Drosophila cellular-ization requires zygotic dunk activity. en. Development 143, 2417–2430 (July 2016).

40. Minestrini, G., Harley, A. S. & Glover, D. M. Localization of Pavarotti-KLP in living Drosophila embryos suggests roles in reorganizing the cortical cytoskeleton during the mitotic cycle. en. Mol. Biol. Cell 14, 4028–4038 (Oct. 2003).

41. Herant, M., Marganski, W. A. & Dembo, M. The mechanics of neutrophils: synthetic modeling of three experiments. en. Biophys. J. 84, 3389–3413 (May 2003).

42. Hibbeler, R. C. Engineering Mechanics: Statics en (Prentice Hall, 2010).

43. Tran, D. T. & Ten Hagen, K. G. Real-time insights into regulated exocytosis. en. J. Cell Sci. 130, 1355–1363 (Apr. 2017).

